# Comparison of evolutionary rescue via biological and cultural evolution

**DOI:** 10.64898/2026.08.27.747706

**Authors:** Shota Shibasaki

## Abstract

Rapid evolution allows populations to persist in environments where they would otherwise go extinct. This phenomenon, known as evolutionary rescue, is typically studied in the framework of biological evolution, yet adaptive traits can also arise and spread through cultural evolution. The present study developed a stochastic eco-evolutionary model to compare rescue probabilities through biological and cultural evolution. Transmission bias governed the rescue probability under cultural evolution by setting how readily a rare adaptive trait was copied. Conformity bias suppressed population persistence because a rare trait was the least likely to be copied. Content bias toward the adaptive trait enabled evolutionary rescue when social learning was rapid, but it typically yielded a lower rescue probability than biological evolution. Only anticonformity bias, together with a high social learning rate, exceeded the rescue probability of biological evolution. These results demonstrate that transmission bias alters the demographic consequences of cultural evolution and highlight the importance of transmission processes in evolutionary rescue theory. Understanding how adaptive behaviours are socially transmitted may also improve predictions of animal population persistence and inform conservation efforts in rapidly changing environments.

## 1 Introduction

Evolutionary dynamics can be fast enough to affect ecological processes (Schoener, 2011). Previous studies have shown that evolutionary responses to abiotic environmental changes (Anstett et al., 2026), competitors (Leonard et al., 2026), or predators (Becks et al., 2010) alter population dynamics. One important consequence for such rapid contemporary evolution is “evolutionary rescue,” in which population extinction is avoided by adaptation (Bell, 2017; Uecker et al., 2026). Laboratory experiments have demonstrated evolutionary rescue in microorganisms (Lindsey et al., 2013; Killeen et al., 2017; Shibasaki and Yamamichi, 2026a) and identified factors that facilitate or hinder it (Carlson et al., 2014; Bell, 2017). Although evolutionary rescue was once considered difficult to observe outside laboratory settings (Carlson et al., 2014), accumulating evidence suggests that it also occurs in the wild in response to environmental change (reviewed in Balstad et al., 2026). This growing body of evidence highlights the need to understand the ecological and evolutionary conditions under which evolutionary rescue is likely to occur. Such knowledge is essential for predicting population persistence and for informing conservation strategies, natural resource management, and agricultural practices (Alexander et al., 2014; Vander Wal et al., 2013; Madgwick et al., 2024).

While most previous studies have focused on evolution through genetic inheritance (hereafter, biological evolution), adaptive traits can also be transmitted through social learning (hereafter, cultural evolution). Recent studies suggest cultural evolution may alter population dynamics in changing environments (Keith and Bull, 2017; Arbon et al., 2025; Brakes et al., 2019, 2025a). Evidence of social learning and culture has been accumulated across taxa (Whiten, 2021; Webster, 2023), and social learning in the wild can spread traits that could affect population dynamics and persistence (e.g., foraging and anti-predator behaviours; see Basava et al., 2025). Although empirical studies on the interplay between population dynamics and cultural evolution are lacking, recent theoretical studies incorporate cultural evolution into eco-evolutionary dynamics. For example, Fogarty and Kandler (2020) demonstrate that populations can avoid extinction through cultural evolution and call the phenomenon “cultural evolutionary rescue.” Using life-history parameters of mammals and birds, Brakes et al. (2025b) show that animal population size can be recovered when social learning transmits adaptive traits. Cultural evolution, therefore, may provide an alternative pathway to evolutionary rescue.

However, two questions remain regarding the role of cultural evolution in evolutionary rescue. First, the effects of transmission biases (Henrich and McElreath, 2003; Hoppitt and Laland, 2013; Kendal et al., 2018) on evolutionary rescue are understudied. Transmission biases shape what is copied and whom individuals learn from, and thus alter cultural evolution and its consequences for population dynamics. Previous cultural eco-evolutionary models (Fogarty and Kandler, 2020; Brakes et al., 2025b) have largely focused on content bias, which refers to the bias toward some cultural traits based on their inherent properties (Stubbersfield, 2022). Frequencies of cultural traits observed by a social learner are also known to alter the probability that traits are socially learned. Conformity bias refers to cases in which a majority trait is copied more often than its frequency would predict, resulting in positive frequency-dependent copying (Boyd and Richerson, 1985). Anticonformity bias, in contrast, makes rarer traits more likely to be copied, resulting in negative frequency-dependent copying (Acerbi and Bentley, 2014). Because evolutionary rescue relies on the establishment of initially rare adaptive traits, these frequency-dependent biases are expected to affect rescue probabilities.

Second, whether and when cultural evolution promotes evolutionary rescue more effectively than biological evolution remains unclear. Previous studies have compared the dynamics of biological and cultural evolution in a variety of contexts (Smolla et al., 2021), yet their relative consequences for population dynamics have not been systematically evaluated (Shibasaki, 2026). Predicting their relative effects on evolutionary rescue is challenging because biological and cultural evolution differ in how adaptive traits arise and spread. Cultural evolution may provide more opportunities for the acquisition of adaptive traits than biological evolution. This is because social learning allows individuals to acquire traits repeatedly throughout life, whereas biological evolution restricts trait acquisition to inheritance at conception. Indeed, Perreault (2012) shows that cultural evolution can be faster than biological evolution. However, cultural evolution does not necessarily favour the spread of traits that enhance individuals’ survival and fertility. Whereas biological evolution is primarily driven by natural selection, cultural evolution may additionally be shaped by transmission biases that are not necessarily aligned with individual fitness (Cavalli-Sforza and Feldman, 1981). As a result, cultural evolution may hinder the spread of adaptive traits. Together, these differences between biological and cultural evolution generate contrasting expectations about the speed and direction of adaptation, leaving it unclear whether cultural evolution should rescue populations more or less effectively than biological evolution.

The present study developed a stochastic eco-evolutionary model to examine how transmission bias alters the rescue probability and whether cultural evolution can rescue populations more effectively than biological evolution. Conformity bias hindered evolutionary rescue because it suppressed the establishment of initially rare adaptive traits. Both content and anticonformity biases promoted evolutionary rescue, but their effectiveness relative to biological evolutionary rescue was contrasting. Anticonformity bias with a high social learning rate rescued populations more frequently than biological evolution did, while content-biased social learning typically resulted in lower rescue probabilities even when the social learning rate was high. These results highlight the importance of transmission bias for evaluating the potential role of socially transmitted behaviours in wildlife conservation and management.

## 2 Methods

The present model implements the stochastic eco-evolutionary dynamics of asexually reproducing haploid organisms using the Gillespie algorithm. The eco-evolutionary model focused on the dynamics following an abrupt environmental change, assuming two distinct traits of the focal organism: a mutant, adaptive trait (*A*) and a wild, maladaptive trait (*M*) to the new environment. This setting reflects the simplest rescue models in population genetics (Gomulkiewicz and Holt, 1995; Orr and Unckless, 2008), which can easily be extended to cultural evolution models (Fogarty and Kandler, 2020). To facilitate the comparison of evolutionary rescue via two evolutionary processes, biological and cultural evolution were implemented within the same stochastic framework.

The two traits are assumed to differ in birth rates (*b*_*A*_ and *b*_*M*_), while sharing the same natural death rate (*d*) and intraspecific competition coefficient (*c*). The adaptive trait is adaptive in the sense that the birth rate is higher than the death rate, while the maladaptive individuals have a smaller birth rate than their death rate (i.e., *b*_*A*_ *>d > b*_*M*_ ≥ 0). The implementation of biological or cultural evolution alters the ways the numbers of adaptive and maladaptive individuals change over time (see below).

In the monomorphic population with trait *X* = *A, M* , the deterministic population dynamics are assumed to follow the standard logistic growth dynamics:

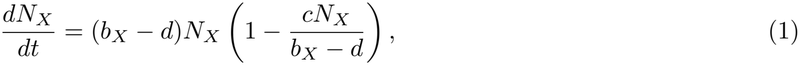

where the intrinsic growth rate is *b*_*X*_ − *d*, and the carrying capacity is (*b*_*X*_ − *d*)*/c*. Because *b*_*M*_ *< d*, the maladaptive population has a negative intrinsic growth rate and declines to zero. In contrast, an adaptive population has a positive intrinsic growth rate *b*_*A*_ − *d >* 0 and approaches its carrying capacity. When the population size (particularly, that of the adaptive individuals) is small, however, evolutionary rescue may fail due to the demographic stochasticity. In the following subsections, the model introduces the stochasticity to investigate how the rescue probabilities change through the biological- and cultural evolution. The analyses of the deterministic models in Supplementary Information provides some necessary conditions for evolutionary rescue (i.e., the presence of an equilibrium with a positive total population size).

### 2.1 Stochastic model without evolution

The first model investigated the stochastic dynamics in the absence of either biological or cultural evolution (Fig. S1). This baseline model clarified that the maladaptive population did not persist without evolution, which is a criterion for evolutionary rescue (Carlson et al., 2014; Uecker et al., 2026). In this scenario, no adaptive individuals existed in the initial population (*N*_*A*_(0) = 0) and never appeared through either mutation, individual learning, or social learning. Then, the birth-and-death reactions are written as follows:

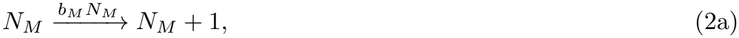

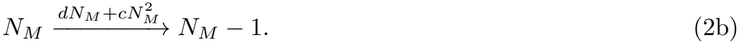

The present study also simulated another baseline model: the dynamics of adaptive population without evolution (Fig. S2). This scenario represents the upper boundary of rescue probability, because population extinction in this scenario indicates the effect of demographic stochasticity after the adaptive trait is fixed in the population.

### 2.2 Biological-evolution model

Next, the model incorporated biological evolution, assuming a one-to-one mapping between traits and genotypes. Although typical models in population genetics analyse the dynamics of alleles or genotypes, the present study uses the term “traits” throughout. This terminology facilitates direct comparison with the cultural evolution models introduced below.

In the presence of biological evolution, an adaptive individual can appear in two ways: when adaptive individuals reproduce offspring without mutation, or when maladaptive individuals reproduce with mutation. Then, the birth-and-death reactions are written as follows (Fig. 1A):

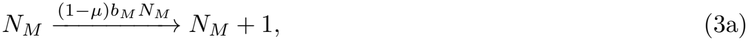

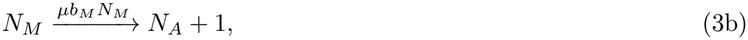

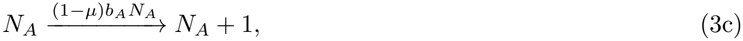

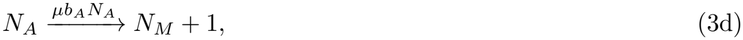

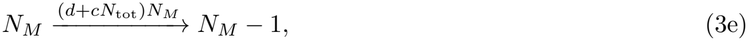

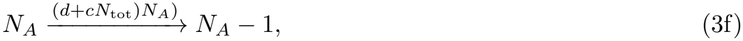

where *µ* is the probability that the mutation occurs. The analytical solution of the rescue probability in Martin et al. (2013) corresponds to the present biological-evolution model with *µ* = 0 and *c* = 0.

**Figure 1:**
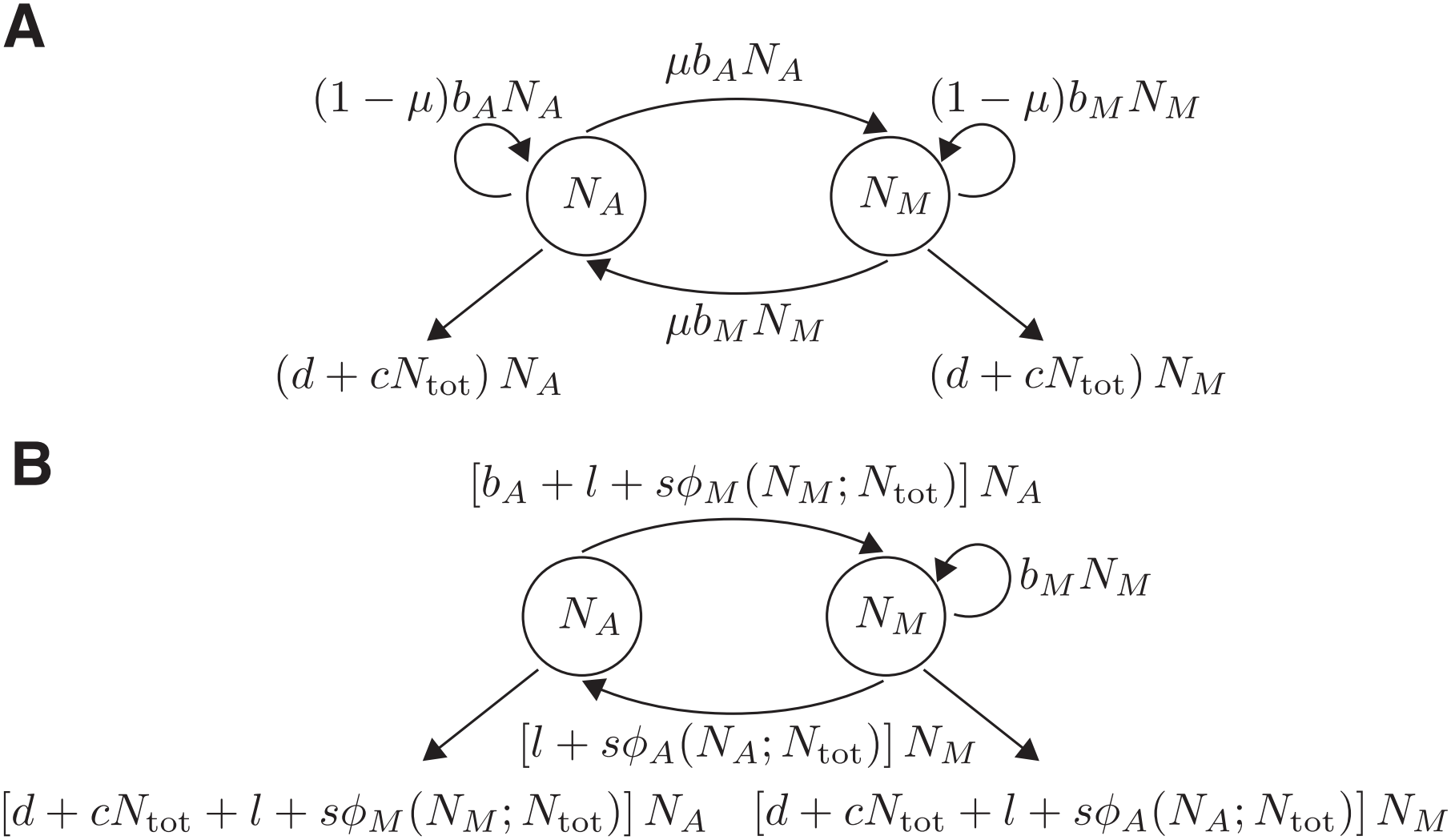
Schematic representation of the eco-evolutionary models. The schematic representations of the eco-evolutionary models are shown. (A): The adaptive individuals appear through biological evolution. (B): The adaptive individuals appear through cultural evolution. Note that the learning processes simultaneously change *N*_*A*_ and *N*_*M*_ while remaining the total population size. The variables and parameters are summarised in Table 1.

### 2.3 Cultural-evolution model

The third model incorporated cultural evolution, in which adaptive traits arose and spread through individual learning and social learning. Individual learning allows individuals to acquire adaptive or maladaptive traits during their lifetime, while social learning enables the transmission of these traits among individuals. Individual learning, therefore, allows reintroduction of traits that have disappeared from the population. This structure parallels the biological-evolution model, where mutation allows genetic variants to be reintroduced after disappearance.

**Table 1:** List of variables and parameters.

| Symbol | Description | Value or range |
|---|---|---|
| *N*<sub>*A*</sub> | Number of adaptive individuals | [0, ∞] |
| *N*<sub>*M*</sub> | Number of maladaptive individuals | [0, ∞] |
| *N*<sub>tot</sub> | Total population size | *N*<sub>*A*</sub> + *N*<sub>*M*</sub> |
| *b*<sub>*A*</sub> | Birth rate of an adaptive individual | [0.11, 0.3] |
| *b*<sub>*M*</sub> | Birth rate of a maladaptive individual | 0.075 |
| *d* | Natural death rate of an individual | 0.1 |
| *c* | Competition coefficient | [10<sup>−4</sup>, 3 × 10<sup>−3</sup>] |
| *μ* | Probability of mutation per birth event | [0, 10<sup>−3</sup>] |
| *l* | Rate of individual learning | [0, 10<sup>−1</sup>] |
| *s* | Rate of social learning | [0, 1] |
| *ϕ*<sub>*i*</sub>(*N*<sub>*i*</sub>; *N*<sub>tot</sub>) | Probability that a social learner copies trait *i* | [0, 1] |
| *w* | Content bias toward the adaptive trait | [−0.5, 1] |
| *θ* | Strength and direction of frequency-dependent bias | [−4, 2] |

The cultural-evolution model differed from the biological-evolution model in three key respects (see also Supplementary Information S1). First, individuals were not born with adaptive traits. All newborns were assumed to be initially maladaptive, and the adaptive trait was acquired only through individual or social learning during an individual’s lifetime, as in previous studies (Thorogood et al., 2017; Kikuchi and Simon, 2026). This contrasted with the biological-evolution model, in which individuals were born adaptive due to genetic inheritance. Second, high individual and social learning rates allowed individuals to change their traits multiple times throughout their lifetimes under cultural evolution, whereas biological evolution was tied to the birth process. This difference was clear through the comparison of the rates at which the number of adaptive individuals increases: *sϕ*_*A*_(*N*_*A*_)*N*_*M*_ +*lN*_*M*_ under cultural evolution, and (1 − *µ*)*b*_*A*_*N*_*A*_ +*µb*_*M*_ *N*_*M*_ under biological evolution. Third, transmission biases modified how the adaptive trait spread within the population through social learning. Depending on the type and strength of the biases, social learning amplified, suppressed, or stabilised adaptive traits. The present model focused on the content bias and frequency-dependent social learning as they were widely investigated in previous studies and were easy to implement. Model-based (e.g., prestige- and kin-based) biases (Henrich and McElreath, 2003; Kendal et al., 2018) were not analysed in the current study because these transmission biases require detailed information of each individual and are not suitable for the Gillespie algorithm.

The birth-and-death reactions under cultural evolution are written as follows (Fig. 1B):

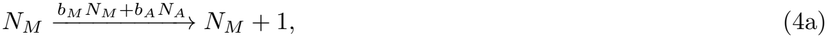

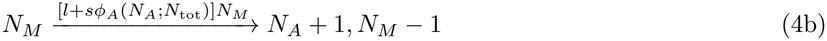

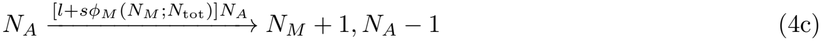

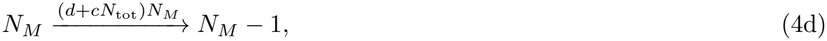

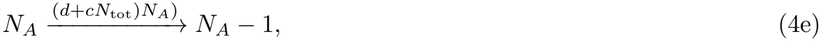

where *l* and *s* are the rates of individual and social learning, respectively, and *ϕ*_*i*_(*N*_*i*_; *N*_tot_) is the probability that a social learner copies the trait *i*.

The probability of socially learning trait *i* varied not only with the number of individuals with adaptive and maladaptive traits, but also with the transmission biases. The present model implemented content-bias and frequency-dependent social learning following the formulation of the previous study (Shibasaki and Yamamichi, 2026b) with a small modification: social learners observed all individuals present in the population at each social learning event (i.e., horizontal, oblique, and vertical transmission were included):

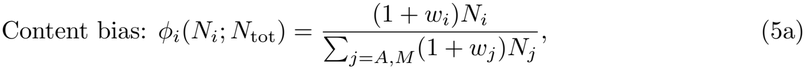

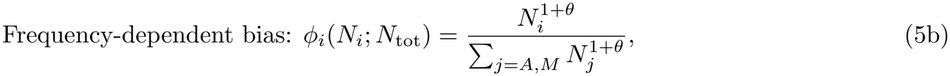

where *w*_*i*_ ≥ −1 represents the weight for trait *i* under content bias, *θ* represents the strength and the direction of frequency-dependent bias. These formulations clarify that the learning probability for trait *i* varies with the type and strength of transmission biases. Without loss of generality, the present study fixed *w*_*M*_ = 0 and used *w* = *w*_*A*_ throughout for simplicity: the weight of the adaptive trait was 1 + *w* while that of the maladaptive trait was 1. In this setting, a positive *w* represents a bias toward the adaptive trait, whereas a negative *w* indicates a bias against it. When *w* = 0 and *θ* = 0, the model returns to unbiased social learning (i.e., random copying). Frequency-dependent bias can be classified into three categories depending on the value of *θ* (Patil et al., 2026): *θ >* 0 corresponds to conformity bias, where a minority cultural trait is adopted by a social learner with a lower probability than under unbiased social learning (Boyd and Richerson, 1985; Efferson et al., 2008). *θ <* 0 corresponds to anticonformity bias, where a minority cultural trait is adopted with a higher probability than under unbiased social learning. Anticonformity bias can be further divided into two sub-categories. Specifically, −1 *< θ <* 0 is referred to as weak anticonformity, because a social learner is more likely to copy the majority cultural trait than the minority trait. In contrast, *θ <* −1 corresponds to strong anticonformity, where the probability of adopting a cultural trait decreases with its frequency. Denton et al. (2020) used alternative formulations to frequency-dependent bias. However, the present formulation is advantageous because it requires only a single parameter, *θ*, regardless of the number of individuals observed during social learning, whereas the number of parameters in the alternative formulation is half the number of observed individuals. If the number of observed individuals (i.e., the number of models) is fixed in the present model as in Denton et al. (2020), the social learning probability cannot be defined when the population size becomes smaller than the number of models.

Note that the present model assumes that social learners cannot copy traits that are absent from the population; once the adaptive trait goes extinct, it is reintroduced into the population only through individual learning. The probability that trait *i* is copied is zero when *N*_*i*_ = 0 in all scenarios. Without this boundary condition, strong anticonformity bias allows a social learner to adopt a cultural trait that does not exist in the population, which violates the definition of social learning.

### 2.4 Simulation setups

The Gillespie algorithm was implemented in Python version 3.11.12. The default values of the birth and death rates of maladaptive individuals were set to *b*_*M*_ = 0.075 and *d* = 0.1, respectively. These parameters determined the timescale of population extinction in the absence of adaptation. Under these default values, maladaptive populations invariably went extinct before *t* = 1000 without any evolutionary processes (Fig. S1). The present study, therefore, defined evolutionary rescue as the persistence of the population until *t* = 1000, i.e., *N*_tot_(1000) = *N*_*A*_(1000) + *N*_*M*_ (1000) *>* 0.

Rescue probabilities were examined across demographic parameters (*b*_*A*_, *c*, and *N*_tot_(0)), evolutionary parameters (*µ, l*, and *s*), and the type and strength of transmission bias (*w* and *θ*; Table 1). The birth rate of adaptive individuals varied as follows: *b*_*A*_ = 0.11, 0.12, 0.15, 0.2, 0.25, 0.3, which enables population persistence when the adaptive trait is fixed (*b*_*A*_ *> d*). The competition coefficient varied as *c* = 10^−4^, 3 × 10^−4^, 10^−3^, 3 × 10^−3^ because the inverse of this parameter affects the carrying capacity of the adaptive population according to equation (1). The carrying capacity of the adaptive population ranged from about three to 2000 with the above demographic parameter values, indicating the variation in the strength of demographic stochasticity. The initial population size was examined in three cases, *N*_tot_ = 10, 100, 1000, while fixing *N*_*A*_(0) = 1, because previous studies have shown that larger initial population sizes can facilitate evolutionary rescue (Bell and Gonzalez, 2009). The evolutionary parameters varied as follows: *µ* = 0, 10^−5^, 10^−4^, 10^−3^, *l* = 0, 10^−5^, 10^−4^, 10^−3^, 10^−1^ , and *s* = 10^−5^, 10^−3^, 10^−1^, 1. The mutation rate was set to represent the rare-mutation regime at the initial condition: *N* (0)*µ ≤* 1. The rates of individual and social learning spanned several orders of magnitude to represent the broad range of successful learning frequencies. When the mortality rate through competition (*cN*_tot_(*t*)) is ignored, *l* = *s* = *d* means that learning and mortality events occur at comparable rates. An individual, therefore, experiences on average *s/d* social learning events during its lifetime when *cN*_tot_(*t*) is small. The upper limit of individual learning was set an order of magnitude below that of social learning, reflecting the assumption that individual learning was harder than social learning. Parameter values for the transmission biases followed the previous study (Shibasaki and Yamamichi, 2026b), except that a wider range of anticonformity strengths was examined: *w* = −0.5, −0.2, −0.1, 0, 0.1, 0.2, 0.5, 1, and *θ* = −4, −3.5, −3, −2.5, −2.1, −1.5, −1.1, −0.9, −0.7, −0.5, −0.3, −0.1, 0.1, 0.5, 1, 1.5, 2.

### 2.5 Statistical analysis

#### 2.5.1 Estimation of rescue probability

For each parameter combination, 1000 independent simulation runs were performed. Let *R* denote the number of rescued populations among the 1000 simulations. The rescue probability *p* was estimated using a Bayesian approach with a uniform prior distribution, *p* ∼ Beta(1, 1). Assuming a binomial likelihood, the posterior distribution of the rescue probability follows Beta(*R* + 1, 1000 − *R* + 1). The rescue probability was summarised as the posterior mean, 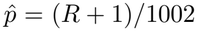, and reported a 95% credible interval (CI) from the 2.5% and 97.5% quantiles of the posterior. This interval reflects the Monte Carlo error arising from the finite number of simulation runs.

#### 2.5.2 Comparison of rescue probability across evolutionary processes

To examine when cultural evolutionary rescue was more likely than biological evolutionary rescue, rescue probabilities between the cultural and biological evolution models were compared. The mutation probability was fixed at maximum (*µ* = 10^−3^) and the individual learning rate was fixed at minimum (*l* = 0). These conditions are conservative because the highest mutation probability maximised rescue probability under biological evolution, whereas the absence of individual learning minimised rescue probability under cultural evolution. Thus, any superiority of cultural evolution observed under these assumptions represents a conservative estimate of the difference in rescue probability between the two evolutionary processes. For each scenario, 20,000 values were sampled from the posterior distributions of rescue probability (see Section 2.5.1) under biological and cultural evolution, and then the posterior distribution of their difference was calculated. The cultural evolution models was compared with biological evolution at each combination of (*b*_*A*_, *c, N*_*M*_ (0)), while varying the type of transmission bias, social learning rate, and the strength of transmission bias within each cultural evolution model.

#### 2.5.3 Measuring mean establishment time

Establishment time was defined as the first time at which the number of adaptive individuals exceeded 10% of the carrying capacity of a population composed entirely of adaptive individuals: *N*_*A*_ *>* 0.1(*b*_*A*_ − *d*)*/c*. This threshold was selected because adaptive individuals reaching this abundance tended to persist (Fig. S2A). Exceeding this threshold did not, however, guarantee long-term persistence, because populations could still go extinct through demographic stochasticity, particularly when carrying capacity was small (i.e., small *b*_*A*_ and *c*; Fig. S2B).

The mean establishment time was conditional on the population persisting at the end of the simulation. However, when the number of successful rescue events was small given the parameter set, the conditional mean establishment time was not reliable. For this reason, the analysis of mean establishment time was restricted to the cases with rescue probability higher than 0.01 (see Table S1 for the number of analysed and removed parameter sets). For each combination of (*b*_*A*_, *c, N*_*M*_ (0)), the difference in mean rescue probability and the difference in mean establishment time (cultural evolution minus biological evolution) were calculated. section Results

### 2.6 Population extinction without evolution

A necessary condition for evolutionary rescue was first confirmed: population extinction in the absence of evolutionary processes (Carlson et al., 2014; Uecker et al., 2026). When the initial population consisted of 1000 maladaptive individuals and the adaptive individuals could not arise through mutation, individual learning, or social learning, the population always went extinct (Fig. S1A). This pattern was consistent over the strength of intraspecific competition and the initial population size (Fig. S1B). These results confirm that any long-term population persistence observed in the following analyses constitutes evolutionary rescue.

Although an adaptive population should persist in the deterministic model, demographic stochasticity led to population extinction even when all individuals were initially adaptive (Fig. S2). In particular, the adaptive population never persisted when (*b*_*A*_, *c*) = (0.11, 0.001), (0.11, 0.003), and (0.12, 0.003) because of its small carrying capacity. In the following evolutionary models, rescue probability remained zero in these cases.

### 2.7 Biological evolution enables evolutionary rescue

Biological evolution enabled evolutionary rescue (Fig. 2; see also Supplementary Information S2.1). Introducing a single adaptive individual to the initial condition rescued populations with some probability even in the absence of mutation (*µ* = 0). Competition decreased the rescue probability by removing the initially rare adaptive individuals and reducing the carrying capacity. This effect was clear when the simulation results were compared with the analytical rescue probability without competition (Martin et al., 2013): the dashed line in Fig. 2A suggests that rescue probability was almost one without competition. In contrast, the rescue probability was below one and decreased while the competition coefficient increased in the current simulation results. The effect of mutation on the rescue probability depended on the initial population size. When the initial population size was small (Fig. S3), the mutation probability did not change the rescue probability, because mutation rarely provides adaptive individuals (i.e., small *b*_*M*_ *µN*_*M*_), and thus the population persistence depends on the survival of the pre-existing adaptive individual. When the initial population size is moderate (Fig. 2) or large (Fig. S4), mutation improves rescue probability. This is because *de novo* adaptive individuals appear at a rate of *b*_*M*_ *µN*_*M*_. When the mutation rate is fixed, however, larger initial population size increases the death rate of adaptive individuals (*d* + *cN*_tot_) and decreases the rescue probability.

**Figure 2:**
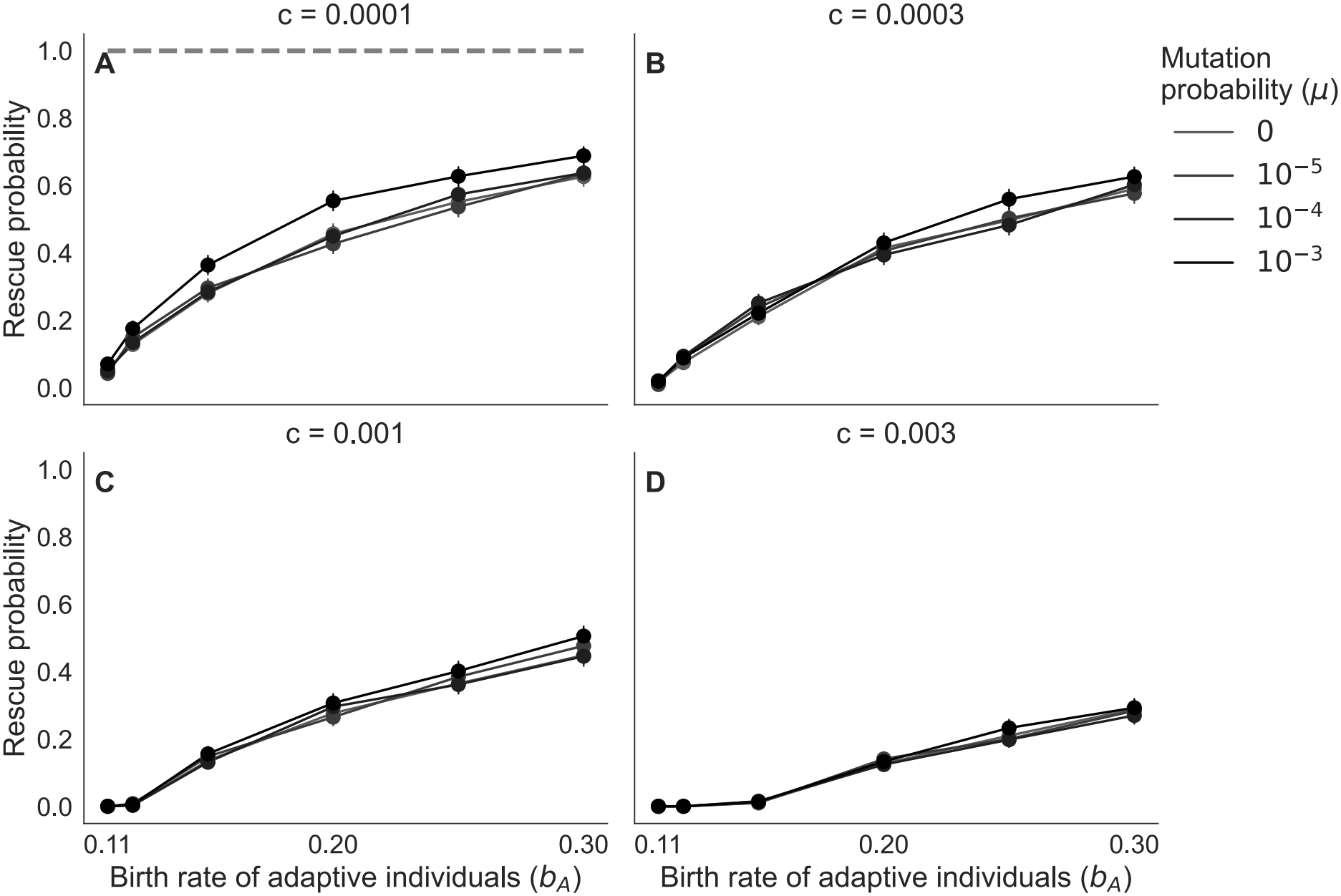
Population can be rescued via biological evolution. The rescue probability under biological evolution is shown over the birth rate of the adaptive individuals *b*_*A*_. The dots represent the proportion of simulations resulting in rescue, and the error bars represent 95% CIs. The dashed line in panel A represents the analytical rescue probability in Martin et al. (2013), which corresponds to the cases of (*c, µ*) = (0, 0) in the present model. The colours correspond to the mutation probability *µ*. Each panel differs in the value of *c*. A: *c* = 0.0001, B:*c* = 0.0003, C: *c* = 0.001, and D: *c* = 0.003. The initial population sizes are *N*_*A*_(0) = 1, and *N*_*M*_ (0) = 99.

### 2.8 Cultural evolution enables evolutionary rescue

The following analysis examined the rescue probability under cultural evolution. In this model, individuals could change their traits without generational turnover, but adaptive traits could be acquired only through individual or social learning. In addition, all newborn individuals were maladaptive in the cultural-evolution model. To clarify the effects of social learning and transmission bias, the present study analysed the simulation results under (i) individual learning alone, (ii) content bias, and (iii) frequency-dependent bias. The analyses of the deterministic models in the Supplementary Information provide necessary conditions for evolutionary rescue (i.e., the presence of an equilibrium with a positive total population size and a non-zero fraction of adaptive individuals). Even if these conditions are satisfied, evolutionary rescue can fail in the stochastic simulations due to demographic stochasticity. Nevertheless, the deterministic model provides the mechanistic understanding of why evolutionary rescue succeeded or failed in the stochastic simulations.

#### 2.8.1 Individual learning alone rarely rescues populations

When the adaptive trait was allowed to arise solely through individual learning (*s* = 0), the population persisted only under a limited range of parameter values. With *l* = 0, 10^−5^, 10^−4^ and 10^−3^, every simulated population went extinct; persistence occurred only at the highest individual learning rate (*l* = 10^−1^; Figs. S5–S7). A lower birth rate of adaptive individuals *b*_*A*_ and a stronger competition coefficient *c* reduced the rescue probabilities when *l* = 10^−1^. The deterministic model explains this pattern (Supplementary Information S2.2). At equilibrium, equation (S8) shows that the frequency of adaptive individuals, *p*^*\**^, increases with *l*, and the total population size, 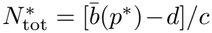, is positive only when (*b*_*A*_ −*b*_*M*_)*p*^*\**^ +*b*_*M*_ *> d*. When *l ≤* 10^−3^, this inequality does not hold for every value of *b*_*A*_ examined. The parameter combinations with 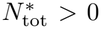 matched those in which rescue occurred with a non-zero probability. The only exception was *l* = 10^−1^ with *b*_*A*_ = 0.15, where 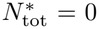. Where 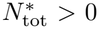, a larger *b*_*A*_ and a smaller *c* increase 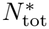, which weakens demographic stochasticity and raises the rescue probability. Individual learning alone, therefore, provided limited opportunities for evolutionary rescue. The following analyses in the main text set *l* = 0 for this reason.

#### 2.8.2 Content bias facilitated cultural evolutionary rescue

Content bias rescued populations even in the absence of individual learning (Fig. 3). However, evolutionary rescue was successful only when the social learning rate was high (*s* = 1) compared to the natural death rate *d* and when the bias favours the adaptive trait (*w >* 0). The deterministic model explains these conditions. An interior equilibrium (0 *< p*^*\**^ *<* 1) exists only when *w >* 0, and the population persistence 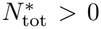 requires *sw > d* (see Supplementary Information S2.3). Rescue through content-biased social learning is therefore possible only when the bias favours the adaptive trait, whereas neither unbiased learning nor a bias towards the maladaptive trait allows the population to persist. The positive effect of the birth rate of adaptive individuals (*b*_*A*_) rapidly saturated. Intraspecific competition (*c*) decreased the rescue probability, as in the biological-evolution model. These patterns were consistent across the initial conditions (Figs. S8 and S9). Introducing individual learning (*l>* 0) resulted in evolutionary rescue with broader ranges of social learning rate *s* and the bias *w*, and enhanced the rescue probability (Figs. S10 and S11).

**Figure 3:**
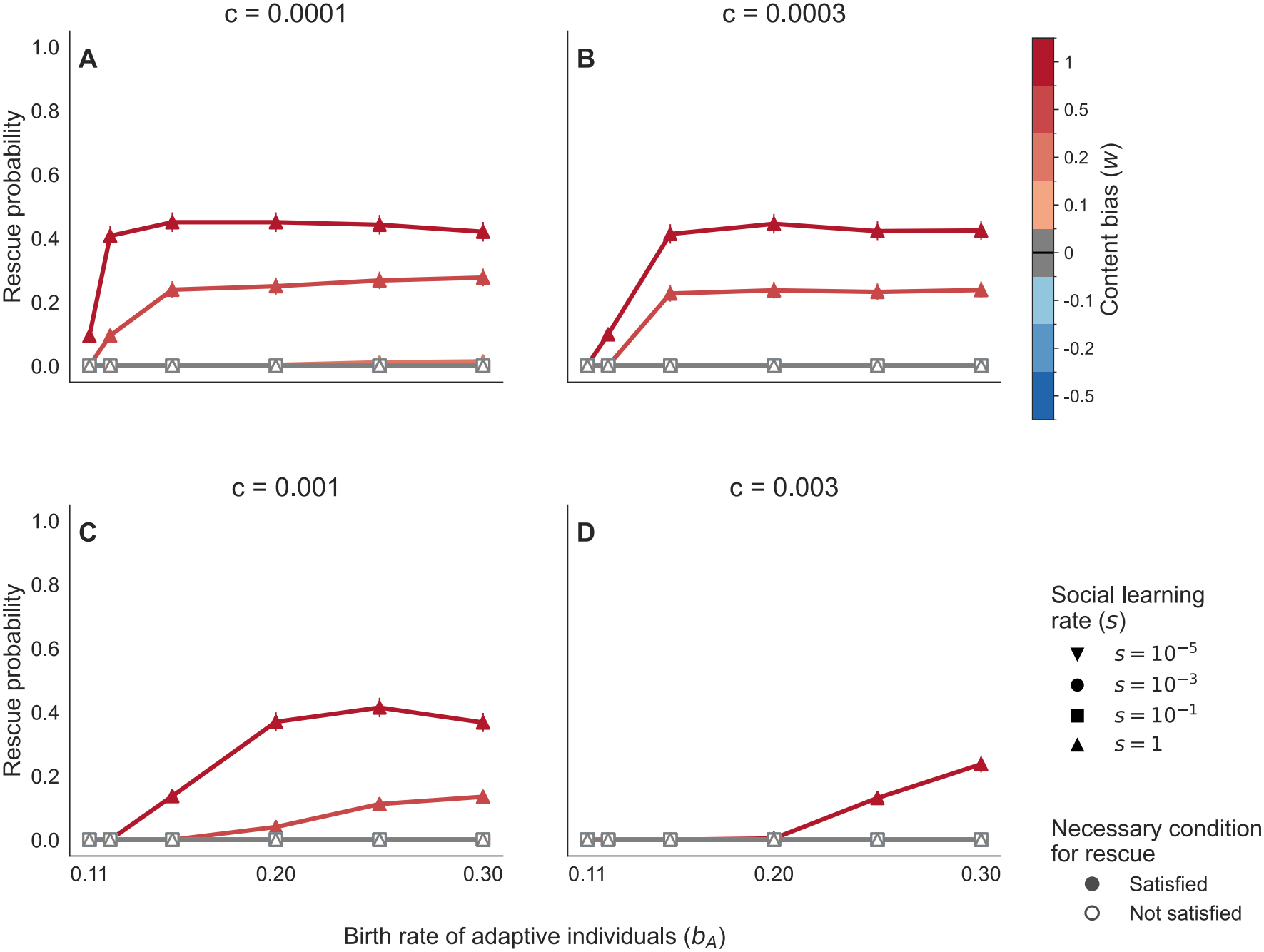
Content bias enables cultural evolutionary rescue. The rescue probability under content-biased cultural evolution is shown as a function of the birth rate of the adaptive individuals, *b*_*A*_. The symbols represent the posterior mean, and the error bars represent 95% CIs. The colours correspond to the bias toward the adaptive trait *w*, and the shapes of the symbols represent the social learning rate *s*. The filled and open symbols represented whether the necessary condition in the deterministic model, inequality (S14), is satisfied or not, respectively. Each panel differs in the value of *c*. A: *c* = 0.0001, B:*c* = 0.0003, C: *c* = 0.001, and D: *c* = 0.003. Parameter values: *l* = 0, *N*_*A*_(0) = 1, and *N*_*M*_ (0) = 99.

#### 2.8.3 Conformity bias impeded cultural evolutionary rescue whereas anticonformity bias facilitated it

Conformity bias (*θ >* 0 in frequency-dependent bias) resulted in population extinction in the absence of individual learning (Figs. 4, S12, and S13). These results indicated that social learning alone was insufficient for evolutionary rescue under this type of transmission bias. Conformity bias favours a majority trait, and thus the initially rare adaptive trait is less often copied than under unbiased social learning (i.e., *w* = 0 in the content bias), which did not rescue the population in the current simulations. However, populations persisted when individual learning rate was high enough to avoid extinction even in the absence of social learning (Fig. S14). In these cases, stronger conformity bias (larger *θ*) and higher social learning rate (larger *s*) decreased the rescue probabilities. Conformity bias, therefore, impeded evolutionary rescue.

**Figure 4:**
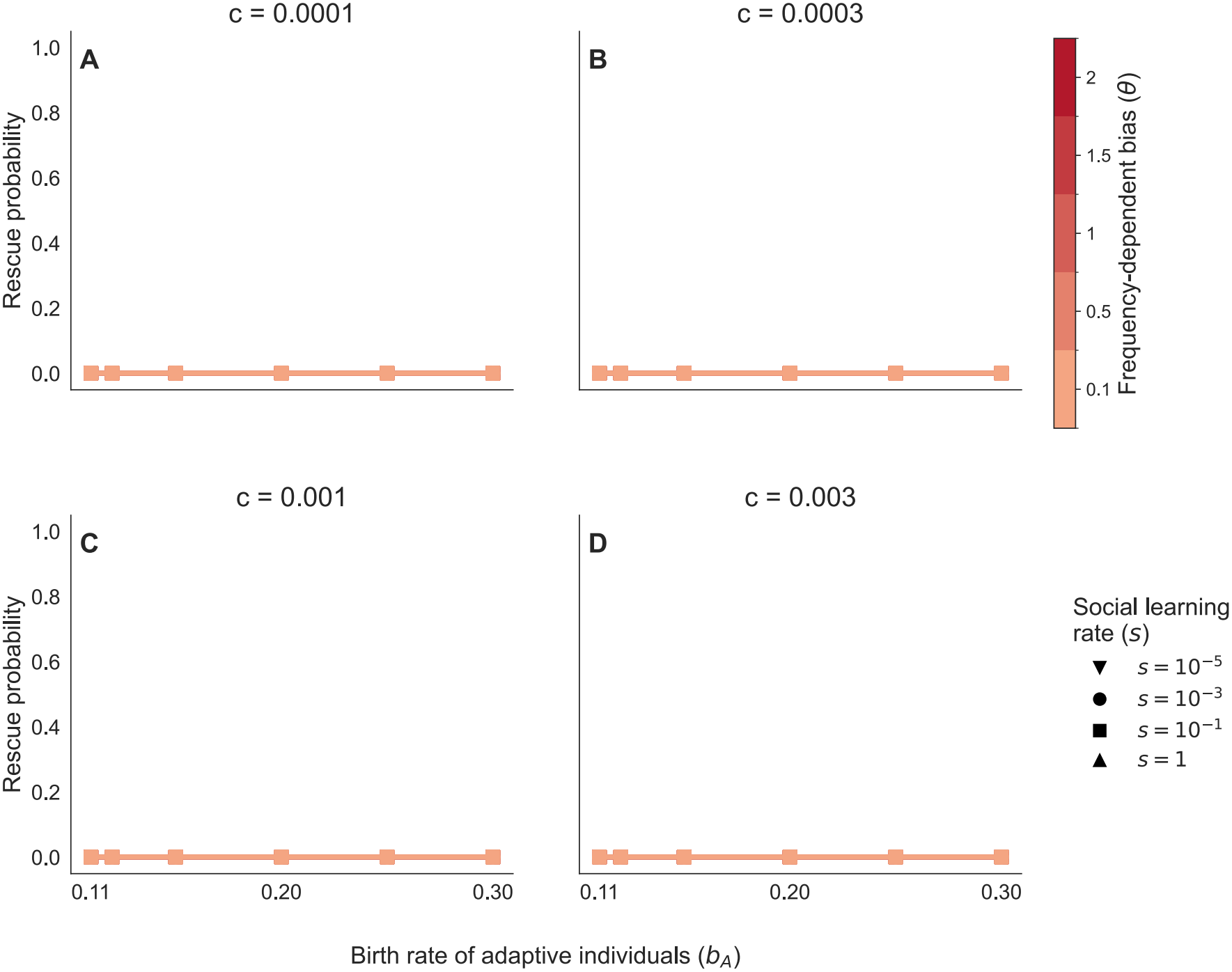
Conformity bias alone could not rescue populations from extinction. The rescue probability under conformity-biased cultural evolution is shown as a function of the birth rate of the adaptive individuals, *b*_*A*_. The symbols represent the posterior mean, and the error bars represent 95% CIs. The colours correspond to the strength of conformity bias *ω*, and the shapes of the symbols represent the social learning rate *s*. Each panel differs in the value of *c*. A: *c* = 0.0001, B:*c* = 0.0003, C: *c* = 0.001, and D: *c* = 0.003. Parameter values: *l* = 0, *N*_*A*_(0) = 1, and *N*_*M*_ (0) = 99.

Anticonformity bias (*θ <* 0 in the frequency-dependent bias) rescued populations even in the absence of individual learning when the social learning rate was high enough (*s* ≥ 0.1; Fig. 5). Although this type of transmission bias has two sub-categories (Patil et al., 2026), the results did not differ qualitatively between them. Stronger anticonformity bias (smaller *θ*) increased the rescue probabilities, because cultural selection favours the initially rare adaptive trait. Furthermore, the initial population size increased the rescue probability (Figs. S15 and S16). Because the current model assumes a single adaptive individual at the beginning of the simulation, increasing the initial population size makes the adaptive trait rarer. According to the learning probability in equation (5b), the rate at which the single adaptive individual is copied, *sN*_*M*_ (0)*ϕ*_*A*_(1) = *sN*_*M*_ (0)*/*(1 + *N*_*M*_ (0)^1+*θ*^), increases with the number of initial population size of the maladaptive individuals *N*_*M*_ (0). The adaptive trait is thus more likely to spread before it is lost when *N*_*M*_ (0) is larger. Unlike content bias, low birth rates of adaptive individuals (*b*_*A*_ ≤ 0.12) did not result in evolutionary rescue. Analysis of the deterministic model suggests that this is because a necessary condition for population persistence at equilibrium is given by the social learning rate and the difference in the birth and death rates of adaptive individuals (See Supplementary Information S2.4 for more details). Anticonformity bias, therefore, facilitated evolutionary rescue only when the birth rate of the adaptive individuals was high enough.

**Figure 5:**
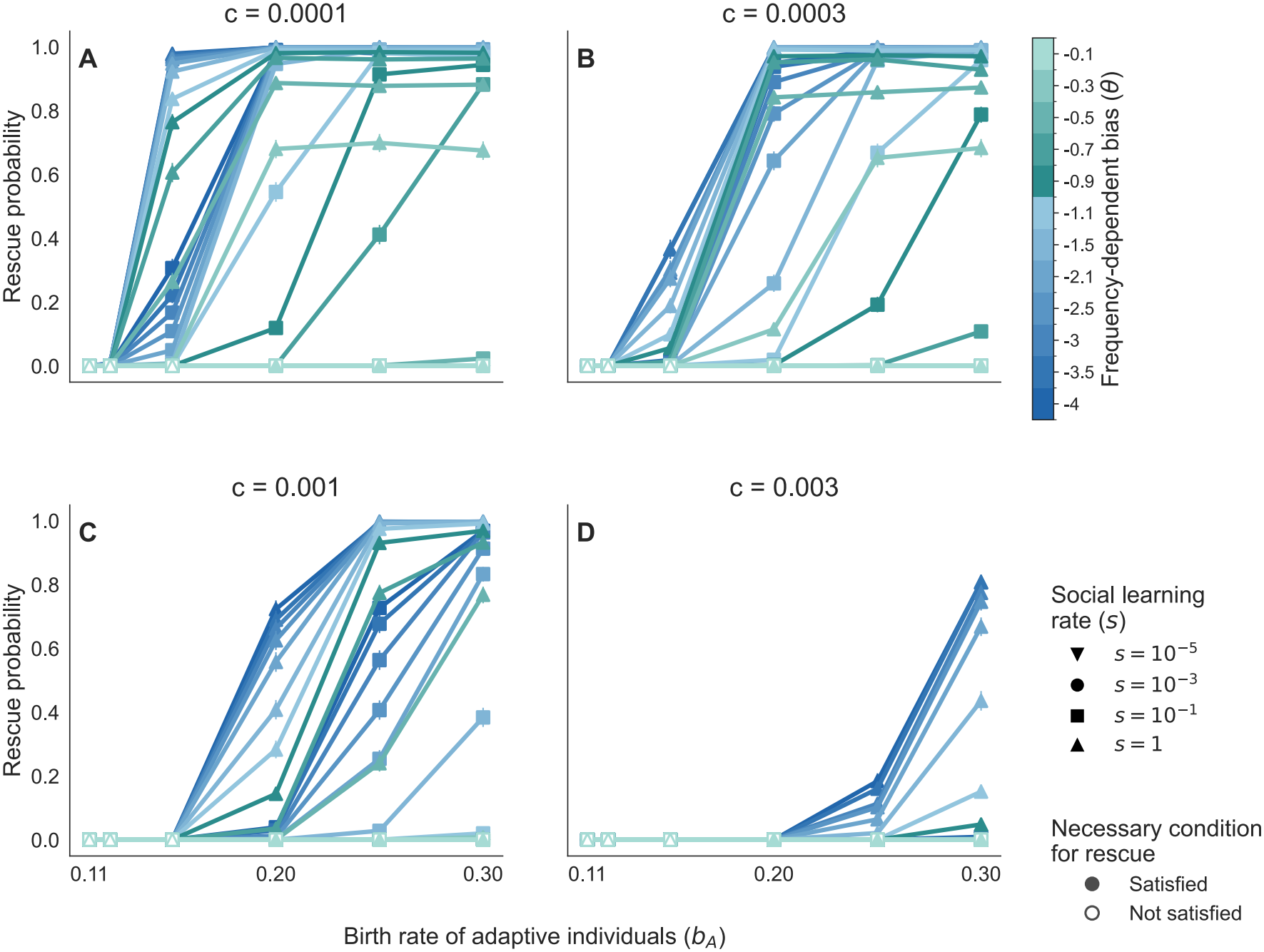
Anticonformity bias promoted cultural evolutionary rescue. The rescue probability under anticonformity-biased cultural evolution is shown as a function of the birth rate of the adaptive individuals, *b*_*A*_. The symbols represent the posterior mean, and the error bars represent 95% CIs. The colours correspond to the strength of anticonformity bias *ω*, and the shapes of the symbols represent the social learning rate *s*. The filled and open symbols represented whether the necessary condition in the deterministic model, inequality (S22), is satisfied or not, respectively. Each panel differs in the value of *c*. A: *c* = 0.0001, B:*c* = 0.0003, C: *c* = 0.001, and D: *c* = 0.003. Parameter values: *l* = 0, *N*_*A*_(0) = 1, and *N*_*M*_ (0) = 99.

Overall, the impacts of social learning on evolutionary rescue depended on the transmission bias. Content and anticonformity biases facilitated evolutionary rescue, whereas conformity bias reduced rescue probabilities. Content and anticonformity biases differed in the parameter space that allowed evolutionary rescue and in the rescue probabilities. Evolutionary rescue occurred at lower social learning rates and was more frequent under anticonformity than under content bias, except when the social learning rate was high (*s* = 1) and the adaptive birth rate was low (*b*_*A*_ ≤ 0.12).

### 2.9 Transmission bias determines whether cultural evolution rescues populations more frequently than biological evolution does

The following analysis compared the rescue probabilities under two evolutionary processes: cultural evolution under conservative conditions (no individual learning) and biological evolution under favourable conditions (the highest mutation rate). Content-biased social learning resulted in a higher rescue probability than biological evolution only when the birth rate of adaptive individuals was low, the social learning rate was high, the bias toward the adaptive phenotype was large, and the competition coefficient was small; otherwise, the difference in rescue probabilities was close to zero or negative (Figs. 6A – D). Conformity bias, on the other hand, resulted in a lower rescue probability than biological evolution (Figs. 6E – H). Anticonformity bias indicated a higher rescue probability than biological evolution when the social learning rate was at least 0.1, and the birth rate of adaptive individuals was high enough (Figs. 6I–L). Smaller initial population size shrank the parameter range where cultural evolutionary rescue was favoured (Figs. S17 and S18), but the qualitative patterns were unchanged. Taken together, transmission bias influenced whether cultural evolution achieved higher rescue probabilities than biological evolution. Whereas conformity bias never increased rescue probability beyond that of biological evolution, content bias and especially anticonformity bias expanded the conditions under which cultural evolution was more likely to rescue populations than biological evolution.

**Figure 6:**
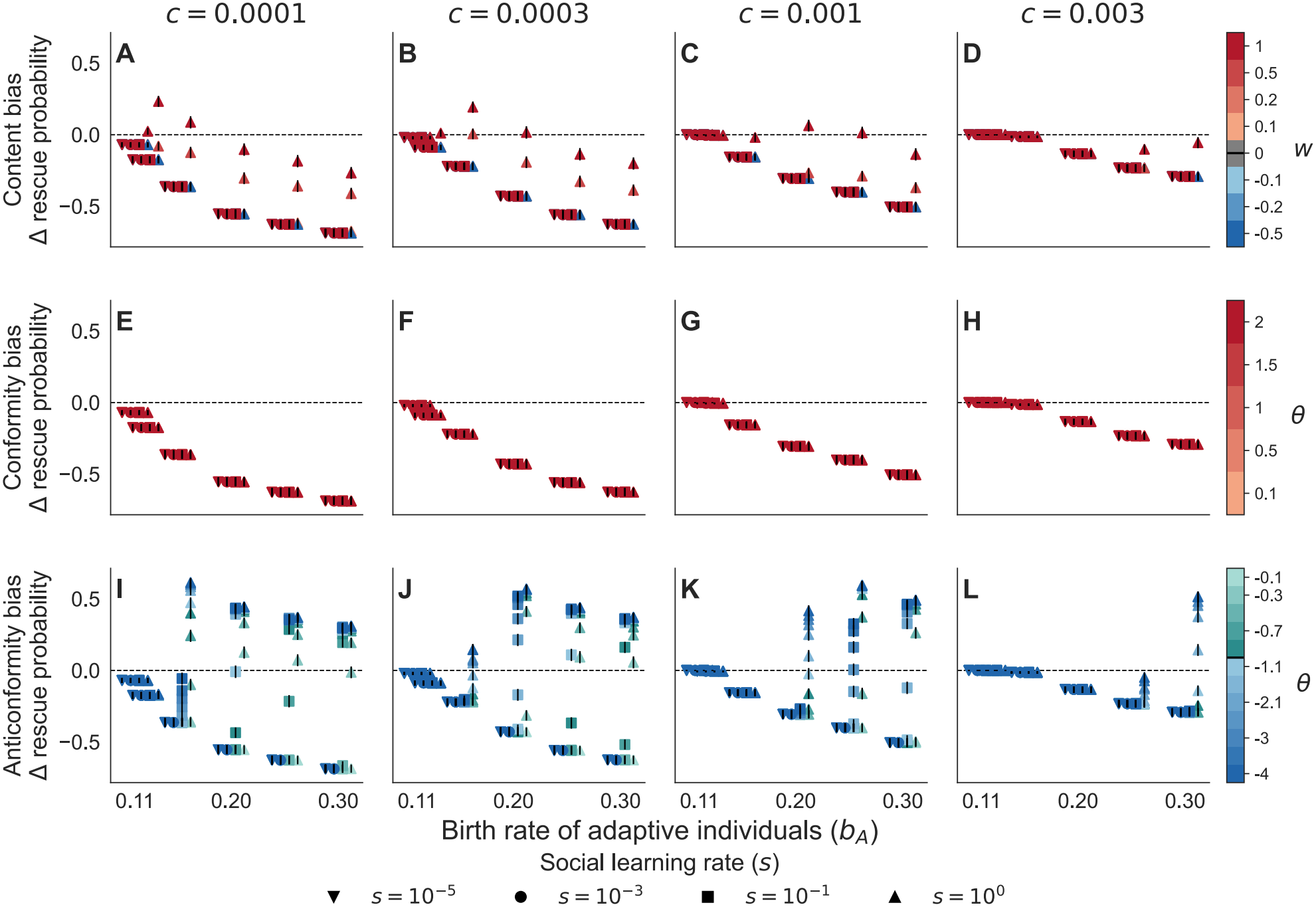
Transmission bias alters the difference in rescue probability between cultural and biological evolution. The rescue probability under cultural evolution minus that under biological evolution is shown as a function of the birth rate of adaptive individuals. Each row differs in transmission bias. A – D: content bias, E – H: conformity bias, and I – L: anticonformity bias. Each column represents the competition coefficient (A, E, and I: *c* = 0.0001; B, F, and J: *c* = 0.0003; C, G, and K: *c* = 0.001; D, H, and L: *c* = 0.003). The symbols represent the mean of the posterior distribution, and the error bars represent 95% CIs. The colours correspond to the strength of bias (*w* and *ω*), and the shapes of the symbols represent the social learning rate *s*. Symbols corresponding to different values of *s* are horizontally offset around each value of *b*_*A*_ to avoid overlap. Parameter values: *µ* = 10^−3^, *l* = 0, *N*_*A*_(0) = 1, and *N*_*M*_ (0) = 99.

Evolutionary rescue requires an initially rare adaptive trait to become established and the population to persist afterwards. A higher rescue probability under cultural evolution might therefore reflect faster establishment of the adaptive trait. When rescue occurred under both biological and cultural evolution (i.e. rescue probability *>* 0.01 in both), however, the difference in mean time to establishment did not generally correspond to the difference in rescue probability (Fig. S19). Cultural evolution both rescued populations more often and established the adaptive trait faster only under anticonformity bias with a large initial population (*N*_tot_ = 100 and 1000). When the initial population was small, establishment under anticonformity bias was often slower than under biological evolution, even when cultural evolution rescued the population more often. Under content bias, the time to establishment was mostly close to that under biological evolution, regardless of the difference in rescue probability. Under conformity bias, the comparison was not possible because cultural evolution did not rescue the population in any parameter set.

## 3 Discussion

Previous studies have demonstrated that populations can avoid extinction through rapid evolution (Balstad et al., 2026; Bell, 2017). While most of these studies have focused on evolutionary rescue through genetic changes, recent work suggests that social learning may offer an alternative pathway to population persistence (Brakes et al., 2025a; Shibasaki, 2026). The present study developed a simple mathematical model to investigate how transmission biases influence cultural evolutionary rescue and when cultural evolution can exceed rescue probabilities under biological evolution. Transmission bias governed whether cultural evolution facilitates or hinders evolutionary rescue. Whereas conformity bias prevented evolutionary rescue (Fig. 4), rescue probabilities were enhanced under content bias toward the adaptive trait and anticonformity-biased transmission (Figs. 3 and 5, respectively) when the social learning rate was high. However, content bias typically resulted in lower rescue probabilities than biological evolution did in a broad parameter space, whereas anticonformity-biased transmission exceeded the rescue probability of biological evolution in a broad parameter space (Fig. 6).

While evolutionary rescue depends on both the emergence of adaptive traits and their establishment before population extinction (Bell, 2017; Uecker et al., 2026), an adaptive individual was already present at the beginning of the simulations. The present study therefore examined the establishment of the adaptive trait under biological and cultural evolution. Natural selection always favours the adaptive trait under biological evolution (see also Supplementary Information S1.1); the fraction of adaptive individuals increases over time, and the rate of its increase through natural selection depends on the difference in the birth rates, *b*_*A*_ − *b*_*M*_ , and the trait variance. Unless demographic stochasticity removes the initially rare adaptive individuals from the population, adaptive traits establish, leading to successful evolutionary rescue. In contrast, under cultural evolution, adaptive traits are transmitted through social learning, and cultural selection (Cavalli-Sforza and Feldman, 1981; Hoppitt and Laland, 2013) acts on the evolutionary dynamics. The rate of change in the fraction of adaptive individuals under cultural selection depends on the social learning rate, *s*, the trait variance, and the transmission biases (see also Supplementary Information S1.2). Moreover, whether cultural selection favours the adaptive trait or not depends on transmission bias, which is independent of the difference in the birth rates of adaptive and maladaptive individuals. In this sense, cultural selection examined in the present study is fitness-blind. Nevertheless, the present study shows fitness-blind social learning rescued the population when the transmission bias facilitated the establishment of an adaptive trait. The bias toward the adaptive trait, *w >* 0, is necessary for evolutionary rescue under content-biased social learning (see Supplementary Information S2.3), and this result is consistent with cultural evolutionary rescue of the previous model (Fogarty and Kandler, 2020; Brakes et al., 2025b). The bias toward the initially rare trait, *θ <* 0, is necessary under frequency-dependent social learning; in other words, conformity bias, *θ >* 0, is maladaptive in the context of evolutionary rescue because it can prevent the spread of the adaptive trait (Eriksson et al., 2007; Kandler and Laland, 2009).

Although the present cultural-evolution model focused on the content-, conformity, and anticonformity-biased social learning, the results provide some insights on how other transmission biases, particularly model-based ones (Kendal et al., 2018), would influence the likelihood of cultural evolutionary rescue. Prestige-based social learning is fitness-blind because individuals with high prestige do not necessarily possess adaptive traits; evolutionary rescue would be successful only if the adaptive trait is associated with prestige. Kin-biased learning (Lamon et al., 2017) is also fitness-blind, because offspring copy their parents regardless of whether the parents carry the adaptive trait. Offspring are still born maladaptive and acquire the trait only through learning. However, because adaptive individuals produce more offspring, more learners would copy adaptive parents, and the birth-rate advantage could indirectly raise the frequency of the adaptive trait. Rescue probabilities under kin-biased learning might approach those under biological evolution. On the other hand, success bias based on reproductive success would be fitness-aligned because social learners tend to copy traits of individuals with more offspring. Based on a previous study (Shibasaki and Yamamichi, 2026b), its dynamics would resemble those under content bias with *w >* 0. Age-based learning would be fitness-aligned if the adaptive trait reduced the death rate, although the current model assumes that the adaptive trait implies a higher birth rate. Under fitness-aligned biases, rescue would be likely when social learning is fast enough. Extending the model to model-based biases would test these predictions.

The deterministic model (Supplementary Information S1) provides necessary conditions for evolutionary rescue, although it ignores demographic stochasticity. The biological model always has an equilibrium with a positive population size, whereas the cultural model has one only when the individual or social learning rate is high enough relative to the death rate (Supplementary Information S2). Whenever these conditions were met, the simulations showed non-zero rescue probabilities (filled symbols in Figs. 3, 5 and S5–S7), except under strong competition and under anticonformity close to unbiased learning. Conversely, rescue occurred in cases in which the equilibrium total population size was zero (individual learning with *l* = 0.1 and *b*_*A*_ = 0.15). This case lies exactly at the threshold, where the population declines very slowly, and rescue became more likely with a larger initial population and weaker competition (Figs. S5–S7).

The deterministic model also helps to explain how the demographic and evolutionary parameters affected rescue probabilities. Unlike in previous studies (Carlson et al., 2014), a larger initial population reduced the rescue probability under biological evolution (Figs. 2, S3 and S4). A larger population intensifies competition, and the single initial adaptive individual must survive population decline, although mutation supplies new adaptive individuals at rate *µb*_*M*_ *N*_*M*_. Under anticonformity bias, by contrast, a larger initial population tended to increase the rescue probability, because a rare trait is adopted by a larger share of learners than its frequency would predict, and a larger population supplies more learners who can copy the single adaptive individual before it is lost. The effect of *b*_*A*_ also differed between the two processes. Under biological evolution, a higher *b*_*A*_ speeds up the spread of the adaptive trait and raised the rescue probability. Under cultural evolution, *b*_*A*_ affects neither cultural selection nor the early spread of a rare adaptive trait; it matters only for whether the population can persist once the trait has spread. The rescue probability therefore saturated with *b*_*A*_ at a level set by the probability that the adaptive trait escapes early loss. When the adaptive trait is rare, content bias raises its chance of being copied almost linearly in *w*; the trait often disappeared before it spread, and the rescue probability saturated below one. Under anticonformity bias, in contrast, a rare trait is copied disproportionately often, and the rescue probability approached one.

Previous studies have suggested that social learning can contribute to population persistence and recovery (Brakes et al., 2019, 2025a), and may therefore serve as a mechanism of evolutionary rescue. Consistent with previous models (Fogarty and Kandler, 2020; Brakes et al., 2025b), the present study supports this view by showing that rapid social learning enabled populations to avoid extinction even when individual learning could not. However, the type of transmission bias altered the likelihood of cultural evolutionary rescue. In particular, rescue probability increased when social learning rates and the strength of these biases were high under content- or anticonformity-biased transmission, whereas conformity bias reversed these relationships (Fig. S14). These findings suggest that the effectiveness of social learning as a conservation tool may depend not only on the presence of social learning itself, but also on the transmission biases through which behaviours spread. Non-human animal culture has been documented across many taxa (Whiten, 2021; Webster, 2023; Basava et al., 2025), and it exhibits a variety of transmission biases (Danchin et al., 2018; van Leeuwen and Hoppitt, 2023). Content and conformity biases are found in animals (Hoppitt and Laland, 2013), but the present analyses suggest these transmission biases are unlikely to yield higher rescue probabilities than biological evolution. In contrast, anticonformity bias can exceed the rescue probability of biological evolution, but clear examples in animal populations are limited (Barrett et al., 2017). These biases are, however, fitness-blind: anticonformity, in particular, would equally spread rare maladaptive traits, which could drive a population to extinction. Future studies are encouraged to investigate the transmission biases used by target animal populations to improve predictions about whether, and under what conditions, cultural evolution can facilitate population persistence.

Several limitations should be acknowledged in this study. The most important limitation is the assumption that adaptive traits can evolve through both genetic changes and social learning. While some behavioural traits may evolve through both biological and cultural evolution, other traits may evolve only through one of these processes. Consequently, the two evolutionary processes may rescue populations by evolving different traits under similar environmental conditions. For example, environmental change that requires access to novel resources can be overcome both through metabolic adaptation in biological evolution (Joshi and Thompson, 1997) and through foraging innovation in cultural evolution (Allen et al., 2013). As a result, comparisons between biological and cultural evolutionary rescue may depend not only on the rate of adaptation but also on the type of adaptive traits available to each evolutionary process. Future studies should investigate scenarios in which biological and cultural evolution generate different adaptive responses to environmental change. Second, this study focused on evolutionary rescue after abrupt changes. Previous studies have also examined rapid evolution in deteriorating and fluctuating environments (Bell, 2017). Evolutionary rescue in these environments typically focuses on the evolution of quantitative traits (Bürger and Lynch, 1995; Peniston et al., 2021; Shibasaki and Yamamichi, 2026a). Future studies can extend the present model to incorporate quantitative traits to compare biological and cultural evolutionary rescue. This research direction would enrich the understanding of the interplay between cumulative cultural evolution and demography (Henrich, 2004; Ghirlanda et al., 2010). A further limitation concerns the parameter dependence of the rescue-probability comparison: The mutation probability *µ* increased rescue probabilities under biological evolution, while the individual learning rate *l* increased them under all three transmission biases. This is why fixing *µ* at its maximum and *l* at its minimum makes the comparison of rescue probability conservative (Fig. 6). Changing these parameter values, therefore, shifts the boundary at which cultural evolution exceeds the rescue probabilities under biological evolution. Nevertheless, the effects of transmission bias on evolutionary rescue remain unchanged.

In summary, the present stochastic eco-evolutionary models demonstrate that transmission bias governs whether cultural evolution facilitates or hinders evolutionary rescue. Whereas conformity bias reduced rescue probability, content and anticonformity biases promoted rescue when the social learning rate was high. When compared with rescue probabilities under biological evolution, anticonformity-biased transmission generated higher rescue probabilities than biological evolution in a broad parameter space, whereas content-biased transmission did so in a limited parameter space. These findings highlight the importance of transmission bias for understanding the ecological consequences of cultural evolution. More broadly, these results suggest that predicting evolutionary rescue requires understanding not only how adaptive traits affect fitness, but also how they are transmitted. Understanding transmission biases may therefore help identify when cultural evolution islikely to contribute to population persistence under environmental change.

## Ethics statement

Not applicable

## Data availability

All simulation data and codes are available from https://doi.org/10.5281/zenodo.24553427.

## Author contributions

Shota Shibasaki conceptualised the project, analysed the model, wrote the original draft, and edited the manuscript.

## Declaration of AI Use

The author used Copilot (Microsoft) and Claude Opus 5.5 (Anthropic) solely to improve the clarity, grammar, and readability of the manuscript. All scientific ideas, mathematical models, analyses, interpretations, and conclusions were conceived and developed solely by the author. All AI-assisted edits were carefully reviewed and verified by the author, who takes full responsibility for the content of the manuscript.

