## Supplementary Information for "Comparison of evolutionary rescue via biological and cultural evolution"

**S3 Supplementary figure for no-evolution model** **7**

**S4 Supplementary figures for biological-evolution model** **9**

**S5 Supplementary figures for individual-learning model** **11**

**S6 Supplementary figures for content-biased social learning model** **14**

**S7 Supplementary figures for conformity-biased social learning model** **18**

**S8 Supplementary figures for anticonformity-biased social learning model** **21**

**S9 Supplementary figures for comparison of rescue probabilities** **23**

**S10 Supplementary figure and table for rescue probability and establishment time** **25**

#### S1 Supplementary text: deterministic eco-evolutionary models

This section derives the deterministic eco-evolutionary models under biological and cultural evolution. Although the deterministic scenarios overlook the effects of demographic stochasticity on evolutionary rescue, these models highlight the key differences in the evolutionary processes, which are important for rescue probability in the main text.

#### S1.1 Eco-evolutionary model under biological evolution

The deterministic dynamics of $N_A$ and $N_M$ under biological evolution are written as follows:

$$\frac{dN_A}{dt} = b_A(1-\mu)N_A + b_M\mu N_M - dN_A - c(N_A+N_M)N_A, \tag{S1a}$$
$$\frac{dN_M}{dt} = b_M(1-\mu)N_M + b_A\mu N_A - dN_M - c(N_A+N_M)N_M. \tag{S1b}$$

These dynamics can be transformed into the dynamics of the total population size $N_A + N_M = N_{\mathrm{tot}}$ (ecological dynamics) and the dynamics of the frequency of adaptive individuals $N_A/N_{\mathrm{tot}} = p$ (evolutionary dynamics).

$$\begin{aligned}\frac{dN_{\mathrm{tot}}}{dt} &= \frac{dN_A}{dt} + \frac{dN_M}{dt}\\ &= \left[\bar{b}(p) - d - cN_{\mathrm{tot}}\right]N_{\mathrm{tot}}, \qquad \text{(S2a)}\\ \frac{dp}{dt} &= \frac{d}{dt}\left(\frac{N_A}{N_{\mathrm{tot}}}\right)\\ &= \frac{1}{N_{\mathrm{tot}}}\frac{dN_A}{dt} - \frac{p}{N_{\mathrm{tot}}}\frac{dN_{\mathrm{tot}}}{dt}\\ &= (b_A - b_M)p(1-p) - \mu\left[b_A p - b_M(1-p)\right], \qquad \text{(S2b)}\end{aligned}$$

where $\bar{b}(p) = pb_A + (1-p)b_M$ represents the mean birth rate. The first term in equation (S2b) represents the effect of natural selection, which is the product of the difference in the birth rate, $b_A - b_M$, and the trait variance $p(1-p)$, while the second term corresponds to the change by mutation. The birth rates $b_A$ and $b_M$ affect these two terms. While the evolutionary processes alter the population dynamics through $\bar{b}(p)$, the population size does not affect the biological evolution.

#### S1.2 Eco-evolutionary model under cultural evolution

The deterministic dynamics of $N_A$ and $N_M$ under cultural evolution are written as follows:

$$\frac{dN_A}{dt} = s\left[\phi_A(N_A;N_{\mathrm{tot}})N_M - \phi_M(N_M;N_{\mathrm{tot}})N_A\right] + l(N_M - N_A) - dN_A - c(N_A+N_M)N_A, \tag{S3a}$$
$$\frac{dN_M}{dt} = \sum_{i=A,M} b_i N_i + s\left[\phi_M(N_M;N_{\mathrm{tot}})N_A - \phi_A(N_A;N_{\mathrm{tot}})N_M\right] + l(N_A - N_M) - dN_M - c(N_A+N_M)N_M. \tag{S3b}$$

Again, these dynamics can be rewritten as the dynamics of $N_{\mathrm{tot}}$ and $p$:

$$\frac{dN_{\text{tot}}}{dt} = \frac{dN_A}{dt} + \frac{dN_M}{dt}$$
$$= \left[\bar{b}(p) - d - cN_{\text{tot}}\right] N_{\text{tot}}, \tag{S4a}$$
$$\frac{dp}{dt} = \frac{d}{dt}\left(\frac{N_A}{N_{\text{tot}}}\right)$$
$$= s \underbrace{\left[(1-p)\phi_A(p_A) - p\phi_M(p_M)\right]}_{g(p)} + l(1-2p) - p\bar{b}(p), \tag{S4b}$$

where the learning probability of trait $i$, $\phi_i(p_i)$, is a function of its frequency $p_i$ (note that $p_A = p$ and $p_M = 1-p$) rather than the number of the individuals. The details of $\phi_i(p_i)$ depends on the transmission bias:

$$\text{Content bias: } \phi_i(p_i) = \frac{(1+w_i)p_i}{\sum_{j=A,M}(1+w_j)p_j}, \tag{S5a}$$
$$\text{frequency-dependent bias: } \phi_i(p_i) = \frac{p_i^{1+\theta}}{\sum_{j=A,M} p_j^{1+\theta}}. \tag{S5b}$$

The first term in equation (S4b) represents the effect of cultural selection, which differs across transmission bias

$$g(p) = \begin{cases} \frac{wp(1-p)}{1+wp} & \text{under content bias,} \\ \frac{p(1-p)[p^\theta - (1-p)^\theta]}{p^{1+\theta}+(1-p)^{1+\theta}} & \text{under frequency-dependent bias.} \end{cases} \tag{S6}$$

Similar to the natural selection term in biological evolution, the cultural selection term includes the trait variance; however, cultural selection is independent of the birth rates. The second term in equation (S4b) shows the impact of individual learning, and the third term indicates the decrease in the frequency of the adaptive trait because newborns are assumed to be born with the maladaptive trait. These formulations clarify that the birth process does not affect cultural evolution through social and individual learning (the first and second terms). As in the biological evolution scenario, the evolutionary processes alter the population dynamics through $\bar{b}(p)$; the population size does not affect the evolutionary dynamics.

Because these deterministic models overlook population extinction through demographic stochasticity, the main text examined the stochastic models and analysed the rescue probabilities across the evolutionary processes and transmission bias.

### S2 Supplementary text: Analyses of the deterministic models

Since $dp/dt$ is independent of $N$ for all cases, $p$ follows a one-dimensional autonomous system and converges either to a boundary ($p^* = 0, 1$) or an interior equilibrium ($0 < p^* < 1$). Then, the total population size converges to $N^*_{\text{tot}}(p^*) = \left[\bar{b}(p^*) - d\right]/c$. If $N^*_{\text{tot}} > 0$, evolutionary rescue is possible, although demographic stochasticity may impede it in the stochastic simulations. Importantly, larger $c$ decreases the total population size in all models examined, and thus it should decrease the rescue probability through demographic stochasticity. If

$N^*_{\text{tot}} < 0$, in contrast, evolutionary rescue is impossible because the population cannot persist in the long term. The presence of an equilibrium with $N_{\text{tot}} > 0$ is, therefore, a necessary condition for evolutionary rescue in the stochastic models. Nevertheless, the analyses of the deterministic models provide mechanistic understanding of why evolutionary rescue succeeded or failed in the main text.

#### S2.1 Biological evolution without mutation

When $\mu = 0$ in the biological-evolution model, $p^* = 1$ and thus $N^*_{\text{tot}}(1) = (b_A - d)/c$ is always positive. However, the simulation results indicate that the rescue probability was almost zero when $b_A$ was small and $c$ was large because the evolutionary rescue failed due to demographic stochasticity (see also Fig. S2B).

#### S2.2 Individual-learning model

When the adaptive individuals arise solely through individual learning ($s = 0$ and $l > 0$ in the cultural-evolution model), an equilibrium is a unique positive root of the following function:

$$h_I(p) = (b_A - b_M)p^2 + (b_M + 2l)p - l, \tag{S7}$$

$$\begin{aligned} h_I(p^*) &= 0, \\ \Leftrightarrow p^* &= \frac{-(b_M + 2l) + \sqrt{(b_M + 2l)^2 + 4l(b_A - b_M)}}{2(b_A - b_M)} \\ &= \frac{2}{\left(\frac{b_M}{l} + 2\right) + \sqrt{\left(\frac{b_M}{l} + 2\right)^2 + \frac{4(b_A - b_M)}{l}}}. \end{aligned} \tag{S8}$$

This equilibrium exists between 0 and 1 because

$$h_I(0) = -l < 0, \tag{S9}$$

$$h_I(1) = b_A + l > 0. \tag{S10}$$

The population can persist if and only if $\bar{b}(p^*) > d$.

#### S2.3 Content-biased social learning model

When adaptive individuals arise only through content-biased social learning ($l = 0$), $p^* = 0$ is always an equilibrium. Dividing $dp/dt = 0$ by $p$ and multiplying by $1 + wp$ shows that any interior equilibrium is a root of

$$h_C(p) = w(b_A - b_M)p^2 + \big[(b_A - b_M) + w(s + b_M)\big]p + (b_M - sw). \tag{S11}$$

When $w = 0$ (unbiased social learning), the only root is $p = -b_M/(b_A - b_M) < 0$; the adaptive trait always goes extinct, and the population cannot persist. When $-1 < w < 0$, no interior equilibrium exists because $h_C(0) = b_M - sw > 0$ and $h_C(1) = (1 + w)b_A > 0$. When $w > 0$, $h_C(p)$ increases with $p$ for $p \geq 0$ and

$h_C(1) = (1+w)b_A > 0$. An interior equilibrium therefore exists, and is unique, if and only if

$$h_C(0) < 0 \Leftrightarrow sw > b_M. \tag{S12}$$

The population then persists if and only if

$$\bar{b}(p^*) = \frac{sw(1-p^*)}{1+wp^*} > d, \tag{S13}$$

whose necessary condition is

$$sw > d \tag{S14}$$

because $(1-p^*)/(1+wp^*) < 1$. Since the simulations in the main text analysed $w \leq 1$, a necessary condition for $N^*_{\text{tot}}(p^*) > 0$ is further simplified as follows:

$$s > d. \tag{S15}$$

#### S2.4 Anticonformity-biased social learning model

Frequency-dependent bias includes both conformity and anticonformity. When the adaptive trait is initially rare, conformity makes it even less likely to be adopted than under unbiased social learning, which itself cannot rescue the population. Rescue under conformity bias is therefore unlikely, and this subsection focuses on the case in which adaptive individuals arise only through anticonformity-biased social learning ($\theta < 0$ and $l = 0$).

Any interior equilibrium frequency of adaptive individuals is a root of

$$s\,G(p) - \bar{b}(p) = 0, \qquad G(p) = \frac{g(p)}{p} = \frac{(1-p)\left[p^{\theta} - (1-p)^{\theta}\right]}{p^{1+\theta} + (1-p)^{1+\theta}}, \tag{S16}$$

where $G(p)$ is the per-capita rate at which social learning increases the frequency of the adaptive trait, divided by the social learning rate $s$. Because of the nonlinearity, the root cannot be obtained in closed form. A necessary condition for $N^*_{\text{tot}} > 0$ can nevertheless be derived as follows.

First, because $b_A > b_M$,

$$N^*_{\text{tot}} > 0 \Leftrightarrow \bar{b}(p^*) > d \Leftrightarrow p^* > \frac{d-b_M}{b_A-b_M} \equiv p_c, \tag{S17}$$

so $p_c$ is the lowest frequency of adaptive individuals at which the population can persist at equilibrium. In addition, $p^* < 0.5$, because $\bar{b}(p^*) > 0$ requires $G(p^*) > 0$, and

$$G(p) \gtreqless 0 \Leftrightarrow p \lesseqgtr 0.5. \tag{S18}$$

The equilibrium frequency therefore satisfies $p_c < p^* < 0.5$, which requires $p_c < 0.5$, that is, $b_A > 2d - b_M$.

Next, $G(p)$ decreases with $p$ on $0 < p < 0.5$. Substituting $r = (1-p)/p$, which satisfies $r > 1$ on this interval, gives

$$\hat{G}(r) = \frac{r - r^{1+\theta}}{1 + r^{1+\theta}}, \qquad \frac{d\hat{G}}{dr} \propto 1 - \theta\, r^{1+\theta} - (1+\theta)\, r^{\theta}. \tag{S19}$$

The second term on the right-hand side is positive because $\theta < 0$. When $\theta \leq -1$, the third term is non-negative, so $d\hat{G}/dr > 0$. When $-1 < \theta < 0$, $r^{\theta} < 1$ for $r > 1$, so $(1+\theta)\, r^{\theta} < 1$ and the sum of the first and third terms is positive; again $d\hat{G}/dr > 0$. Because $dr/dp = -1/p^2 < 0$,

$$\frac{dG}{dp} = \frac{d\hat{G}}{dr}\,\frac{dr}{dp} < 0. \tag{S20}$$

Finally, since $p_c < p^*$ and $G$ is decreasing,

$$s\, G(p_c) > s\, G(p^*) = \bar{b}(p^*) > d. \tag{S21}$$

A necessary condition for $N^*_{\text{tot}} > 0$ is therefore

$$s\, G(p_c) > d, \tag{S22}$$

where

$$G(p_c) = \frac{r_c - r_c^{1+\theta}}{1 + r_c^{1+\theta}}, \tag{S23}$$

$$r_c = \frac{1 - p_c}{p_c} = \frac{b_A - d}{d - b_M}. \tag{S24}$$

### S3 Supplementary figure for no-evolution model

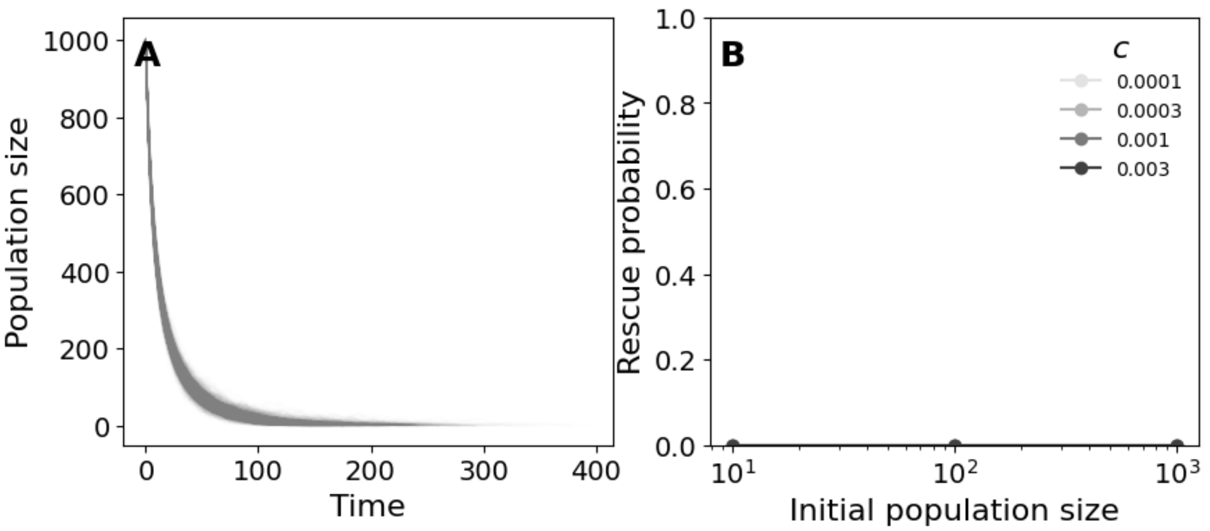

Figure S1: Population extinction in the absence of evolutionary processes

(A) One thousand stochastic population trajectories without mutation, individual learning, or social learning. Parameter values were $c = 10^{-4}$, $N_M(0) = 1000$, and $N_A(0) = 0$. All populations went extinct before $t = 1000$. (B) Rescue probability as a function of the competition coefficient ($c$) and initial maladaptive population size ($N_M(0)$). The initial adaptive population size was fixed at zero ($N_A(0) = 0$). The dots represent the posterior mean, and the error bars represent 95% CIs; they are smaller than the plotting dots.

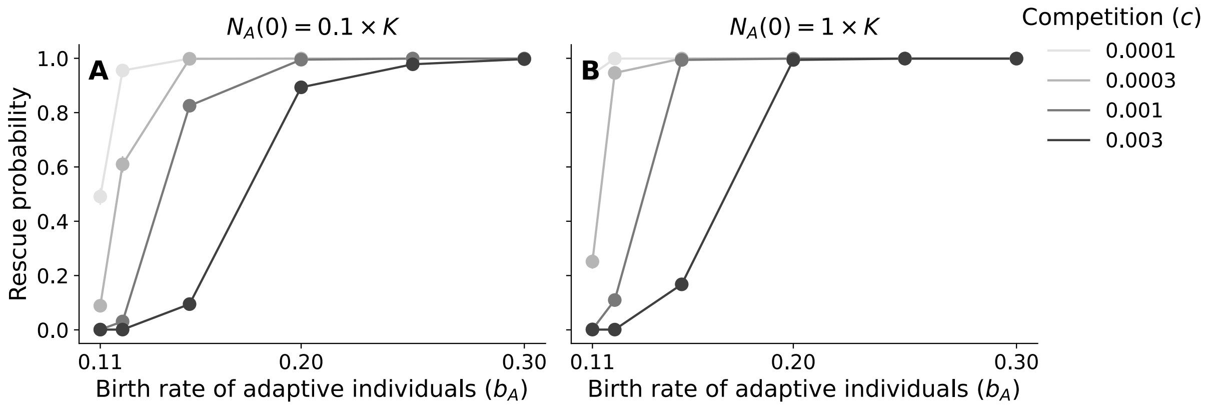

Figure S2: Persistence of adaptive population without evolution

Probability of population persistence as a function of the birth rate of adaptive individuals, $b_A$. (A) Initial population size was set to the floor of 10% of carrying capacity, $\lfloor 0.1(b_A - d)/c \rfloor$. (B) Initial population size was set to the floor of carrying capacity, $\lfloor (b_A - d)/c \rfloor$. No evolution occurred, and maladaptive individuals did not exist at the initial condition ($N_M(0) = 0$).

### S4 Supplementary figures for biological-evolution model

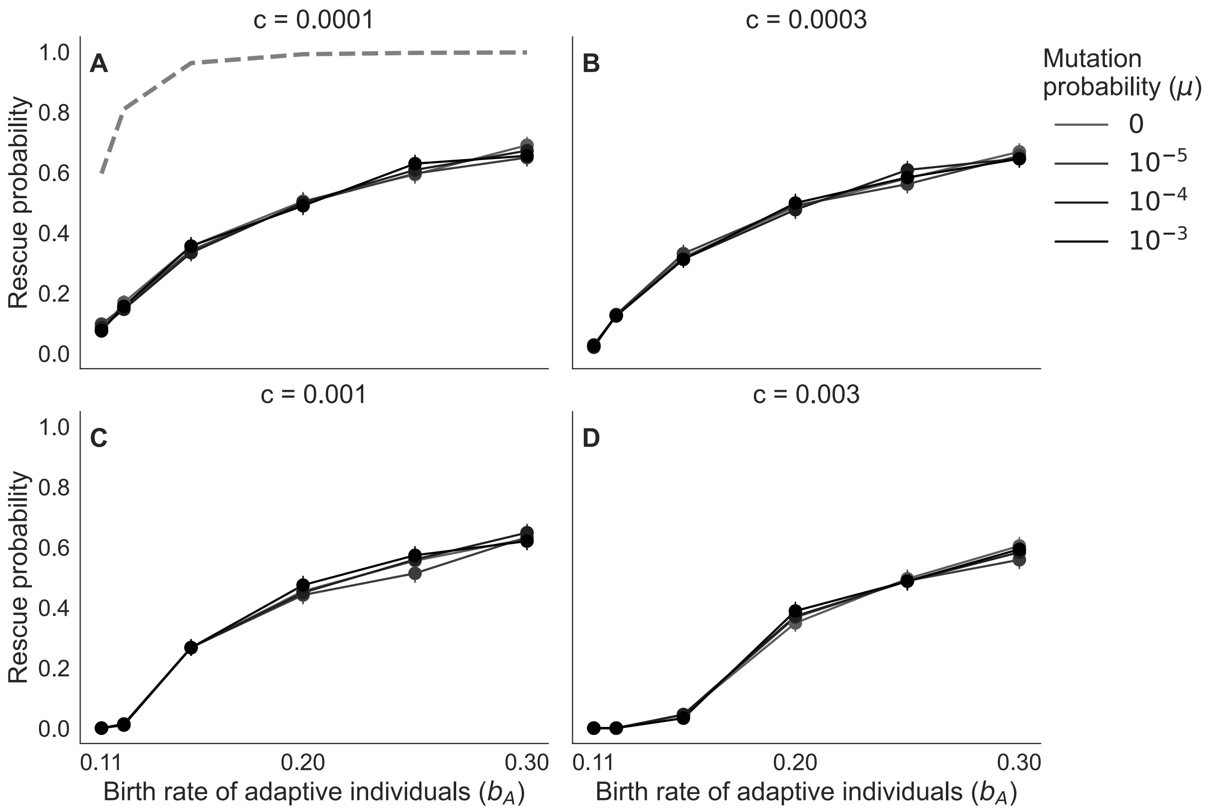

Figure S3: Population can be rescued via biological evolution when the initial population size was smaller

Rescue probabilities in the biological-evolution model are shown with a small initial population size. Similar to Fig. 2, but the initial population sizes are $N_A(0) = 1$ and $N_M(0) = 9$.

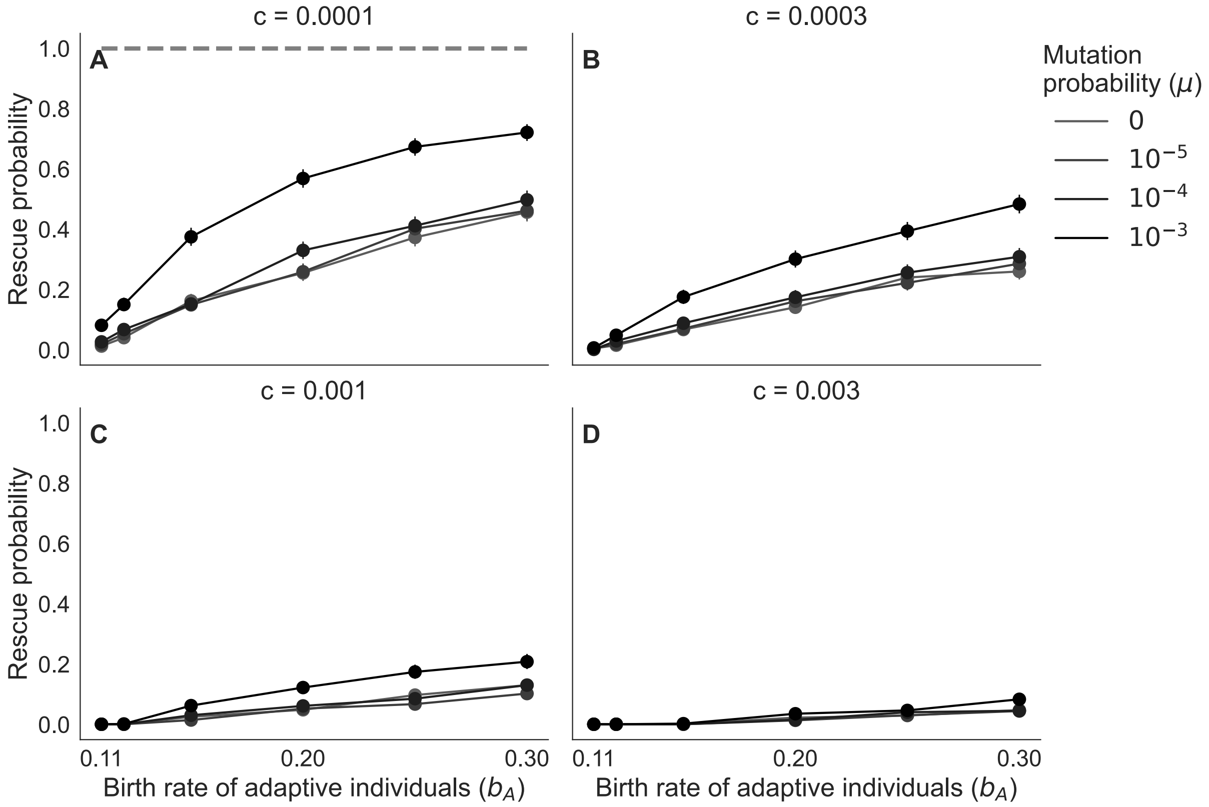

Figure S4: Population can be rescued via biological evolution when the initial population size was larger

Rescue probabilities in the biological-evolution model are shown with a large population size. Similar to Fig. 2, but the initial population sizes are $N_A(0) = 1$ and $N_M(0) = 999$.

### S5 Supplementary figures for individual-learning model

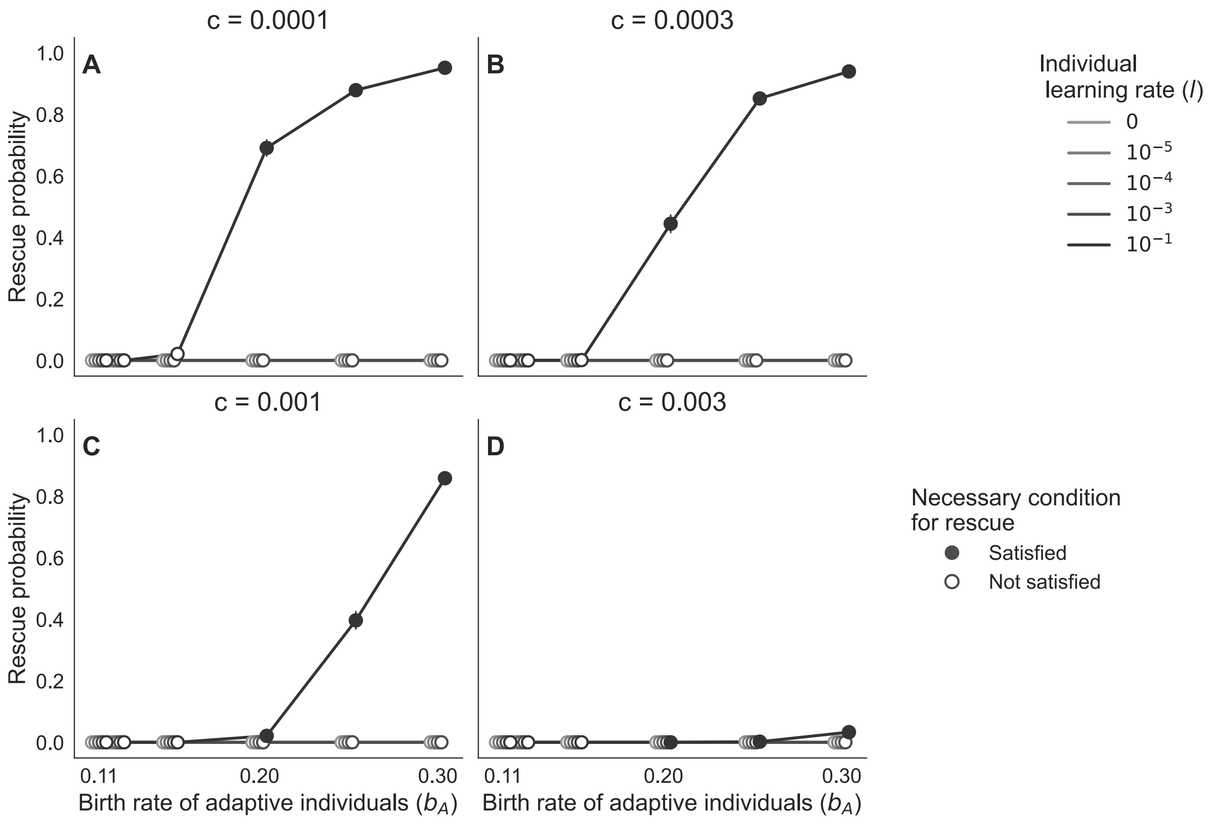

Figure S5: Population can rarely persist solely through individual learning with a small initial population size

Rescue probability under individual learning alone as a function of the birth rate of adaptive individuals ($b_A$). The dots represent the posterior mean, and the error bars represent 95% CIs. Colours represent the individual learning rate ($l$). Panels differ in the strength of intraspecific competition: (A) $c = 0.0001$, (B) $c = 0.0003$, (C) $c = 0.001$, and (D) $c = 0.003$. Filled and open symbols indicate parameter sets where rescue is possible ($N^*_{\text{tot}} > 0$) and impossible ($N^*_{\text{tot}} \leq 0$), respectively, in the deterministic model. Points are slightly offset horizontally for visibility. The initial population sizes are $N_A(0) = 1$ and $N_M(0) = 9$.

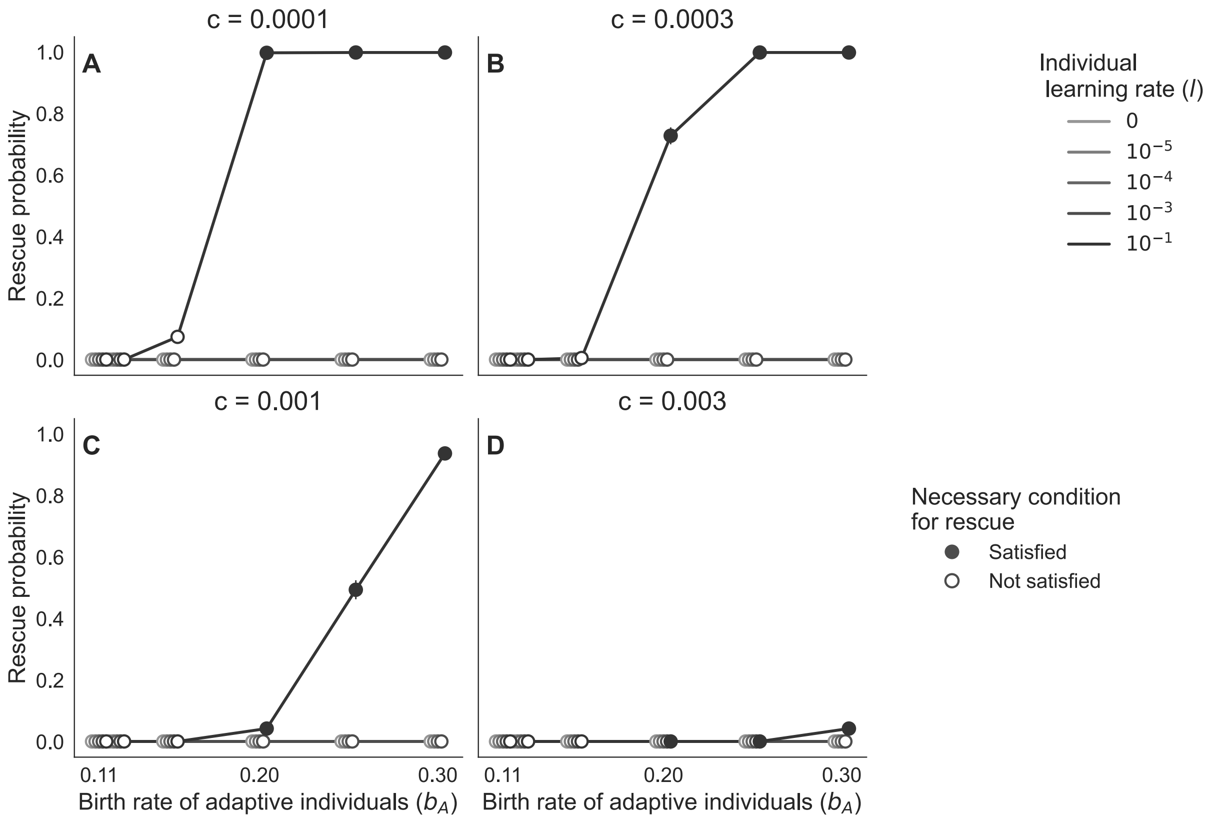

Figure S6: Population can rarely persist solely through individual learning with a moderate initial population size

Similar to Fig. S5, but initial population sizes are $N_A(0) = 1$ and $N_M(0) = 99$.

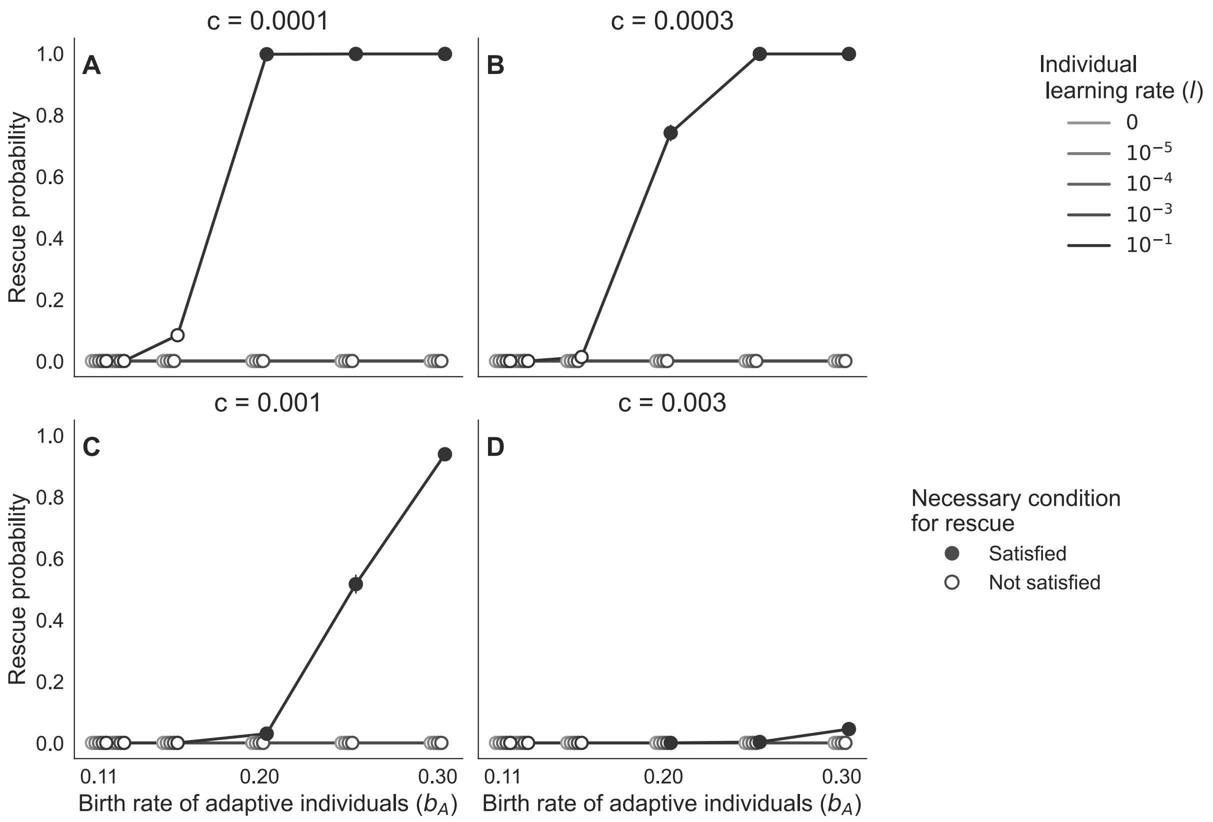

Figure S7: Population can rarely persist solely through individual learning with a large initial population size

Similar to Fig. S5, but initial population sizes are $N_A(0) = 1$ and $N_M(0) = 999$.

### S6 Supplementary figures for content-biased social learning model

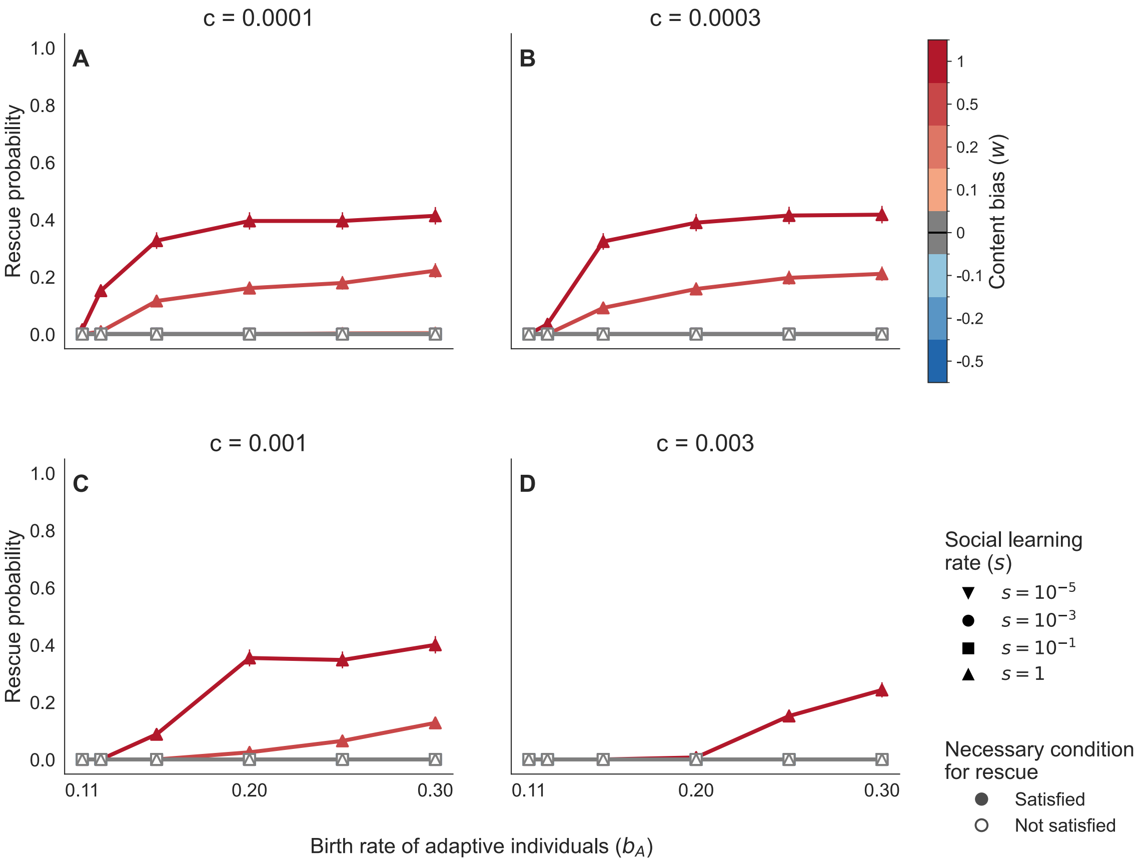

Figure S8: Content bias enables cultural evolutionary rescue with a small initial population size

Similar to Fig. 3, but the initial population sizes are $N_A(0) = 1$ and $N_M(0) = 9$.

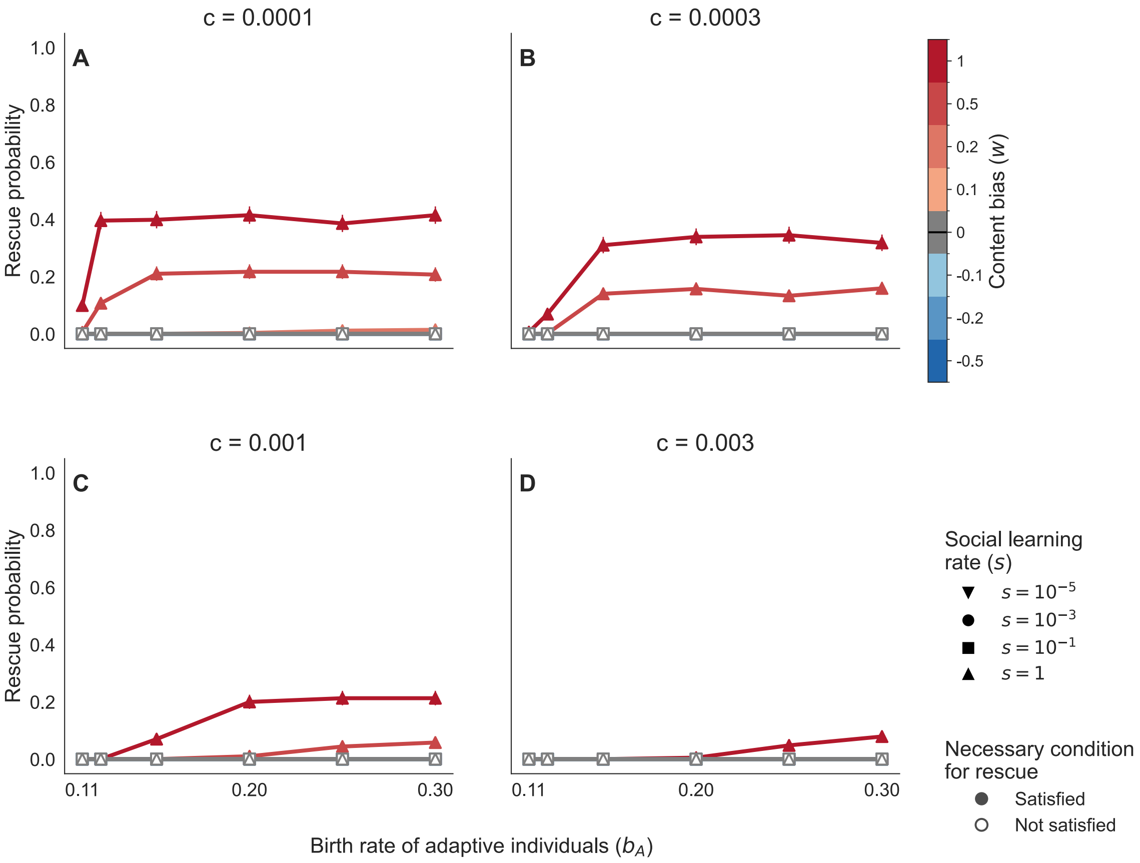

Figure S9: Content bias enables cultural evolutionary rescue with a large initial population size

Similar to Fig. 3, but the initial population sizes are $N_A(0) = 1$ and $N_M(0) = 999$.

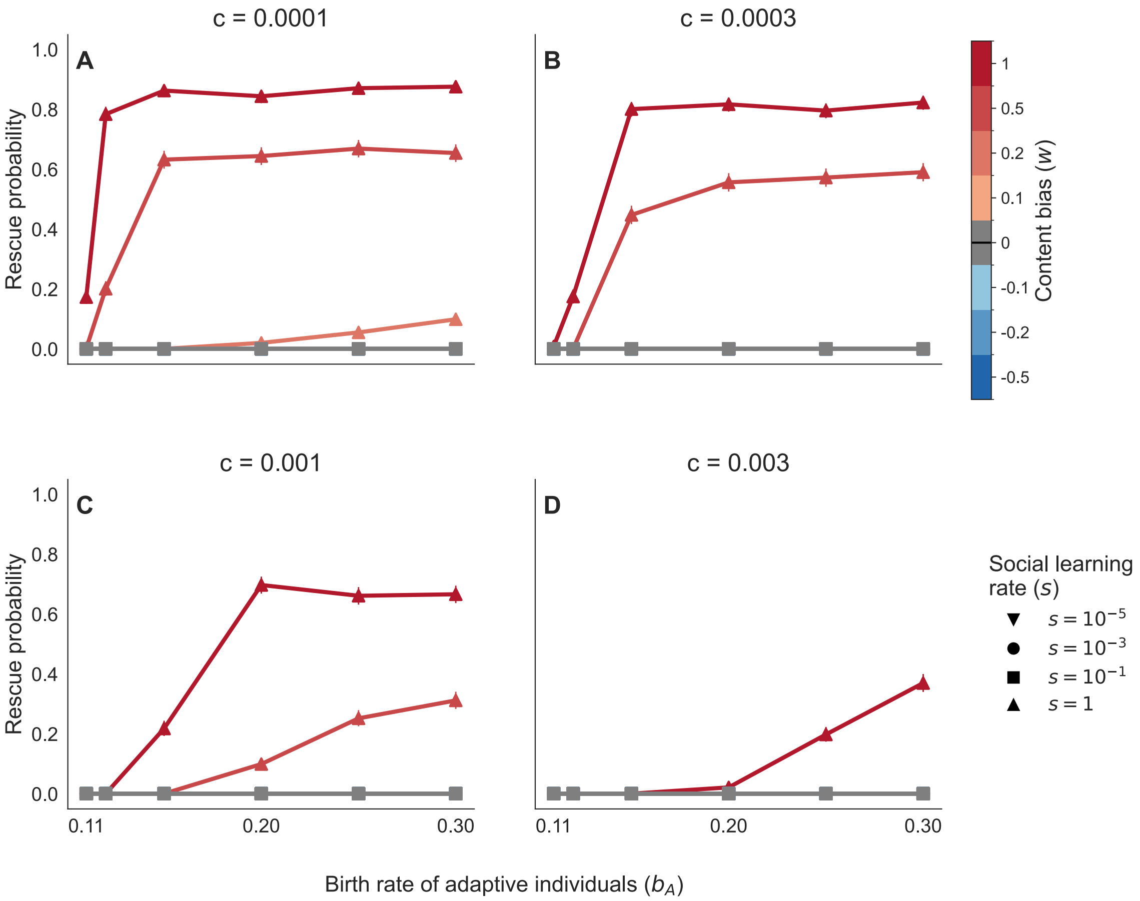

Figure S10: Individual learning enhances rescue probability under content-biased social learning

Similar to Fig. 3, but the individual learning rate is $l = 10^{-3}$. Without social learning, evolutionary rescue never succeed in this case (Fig. S6.)

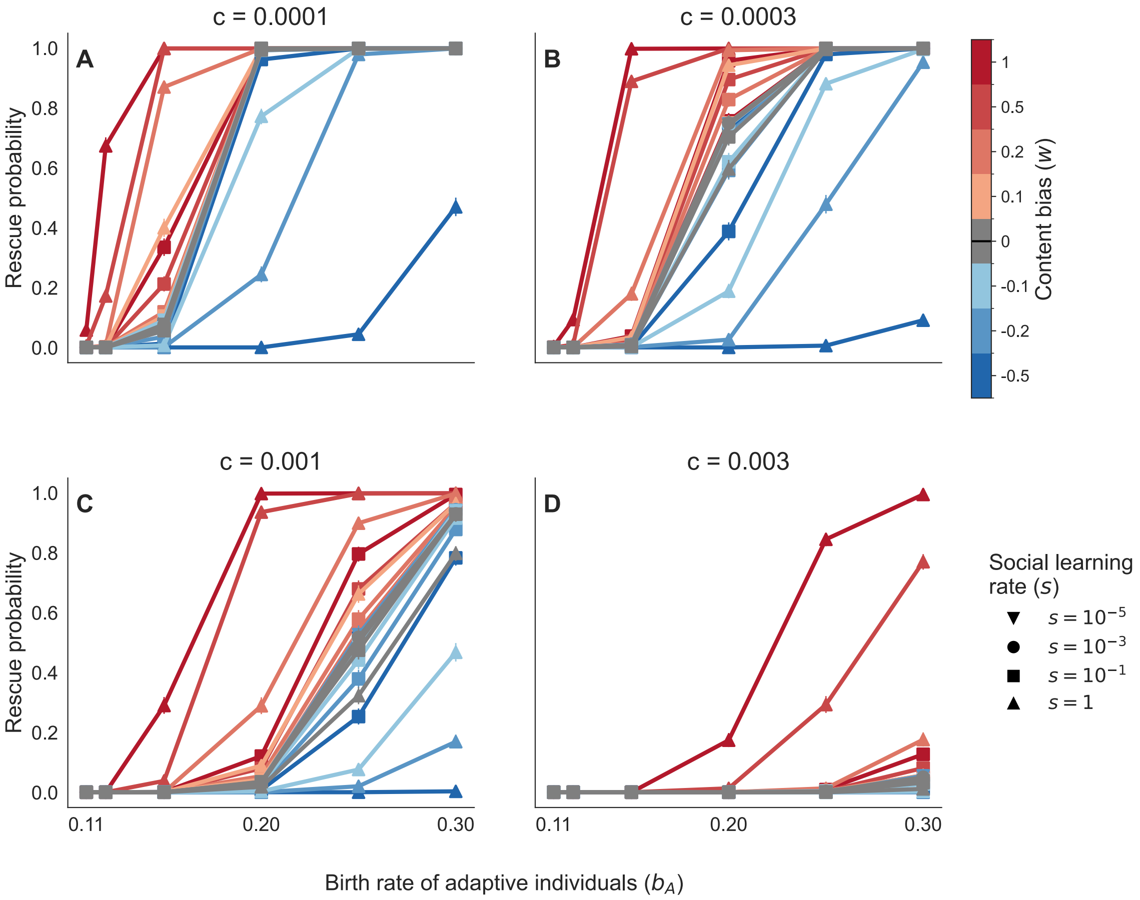

Figure S11: Individual learning allows evolutionary rescue even when content-biased social learning disfavours adaptive traits

Similar to Fig. 3, but the individual learning rate is $l = 10^{-1}$. Because of the high individual learning rate, the population can persist without social learning (Fig. S6)

757 # S7 Supplementary figures for conformity-biased social learning model

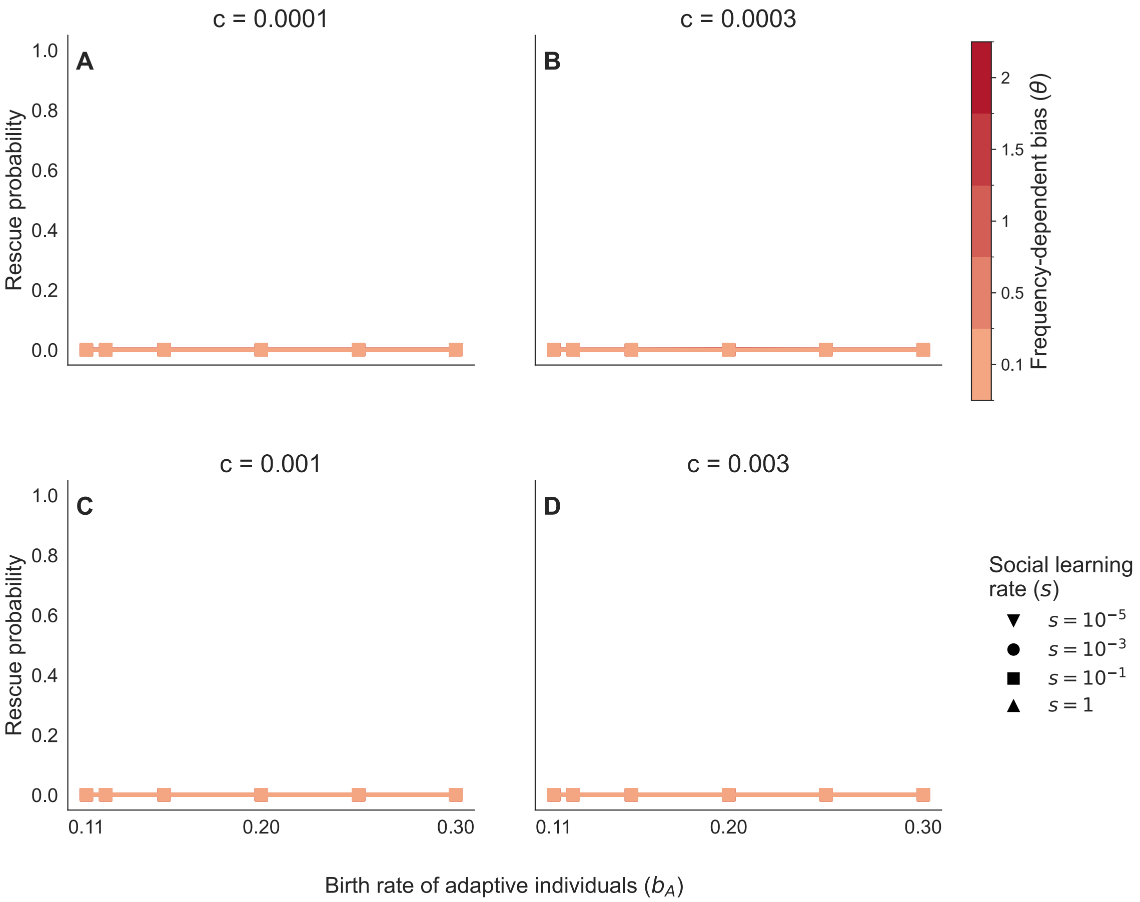

Figure S12: Conformity bias alone could not rescue populations from extinction with a small initial population size

Similar to Fig. 4, but the initial population sizes are $N_A(0) = 1$ and $N_M(0) = 9$.

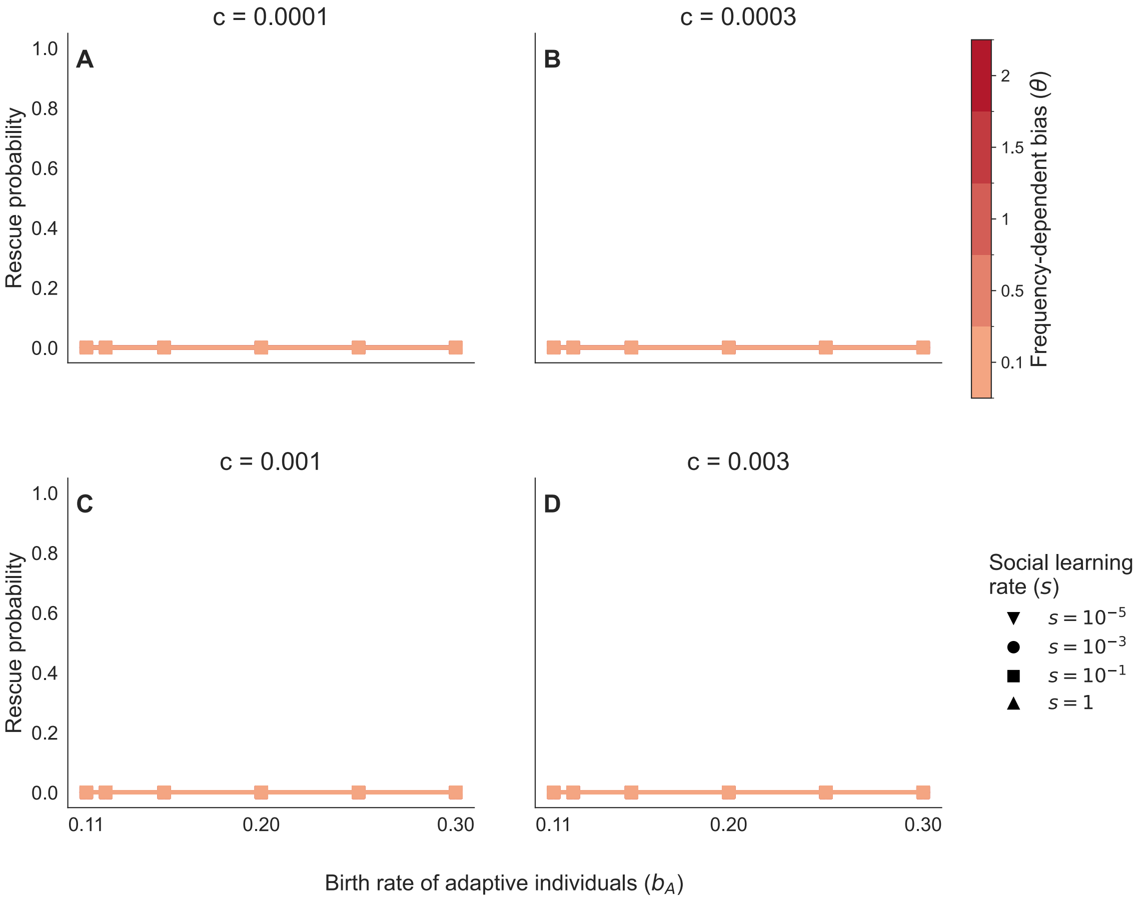

Figure S13: Conformity bias alone could not rescue populations from extinction with a large initial population size

Similar to Fig. 4, but the initial population sizes are $N_A(0) = 1$ and $N_M(0) = 999$.

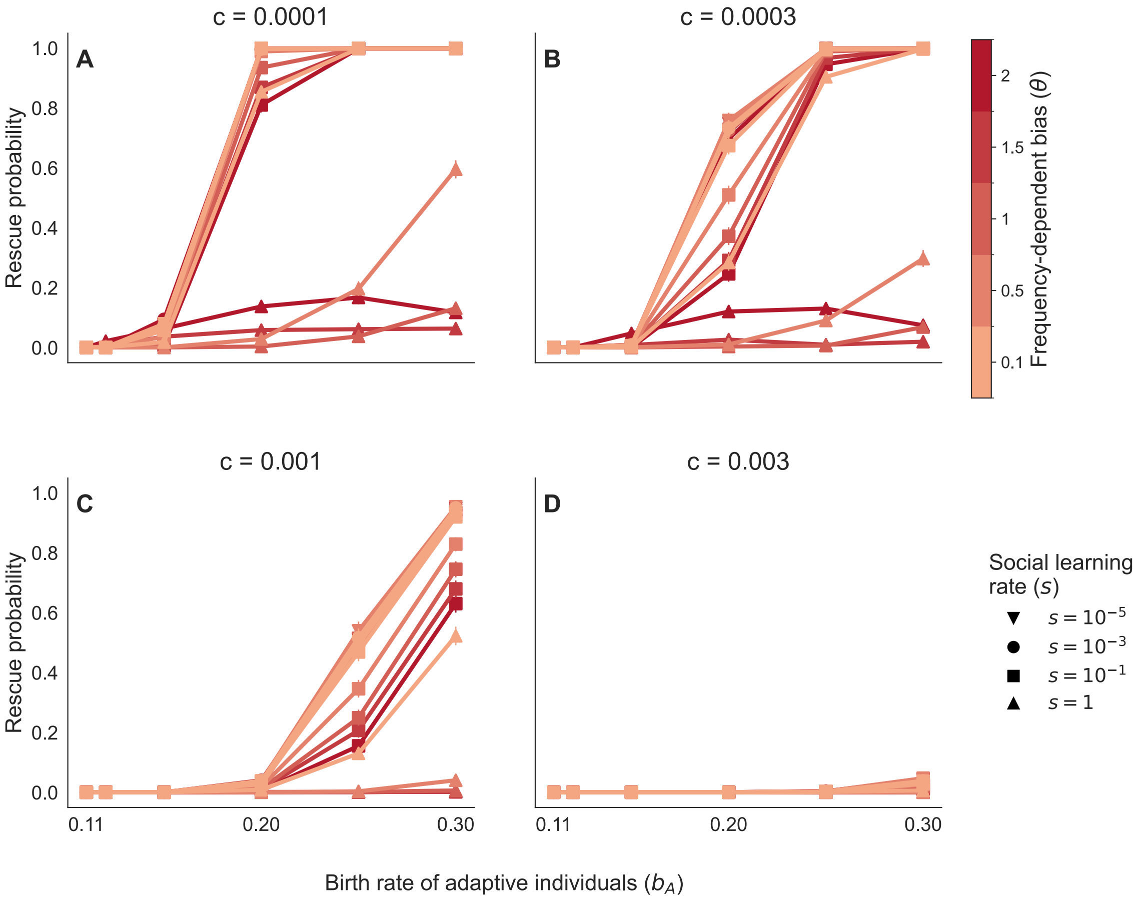

Figure S14: Evolutionary rescue succeeds under conformity-biased social learning if the individual learning rate is sufficiently high

Similar to Fig. 4, but the individual learning rate is $l = 0.1$. Because of the high individual learning rate, the population can persist without social learning (Fig. S6).

### S8 Supplementary figures for anticonformity-biased social learning model

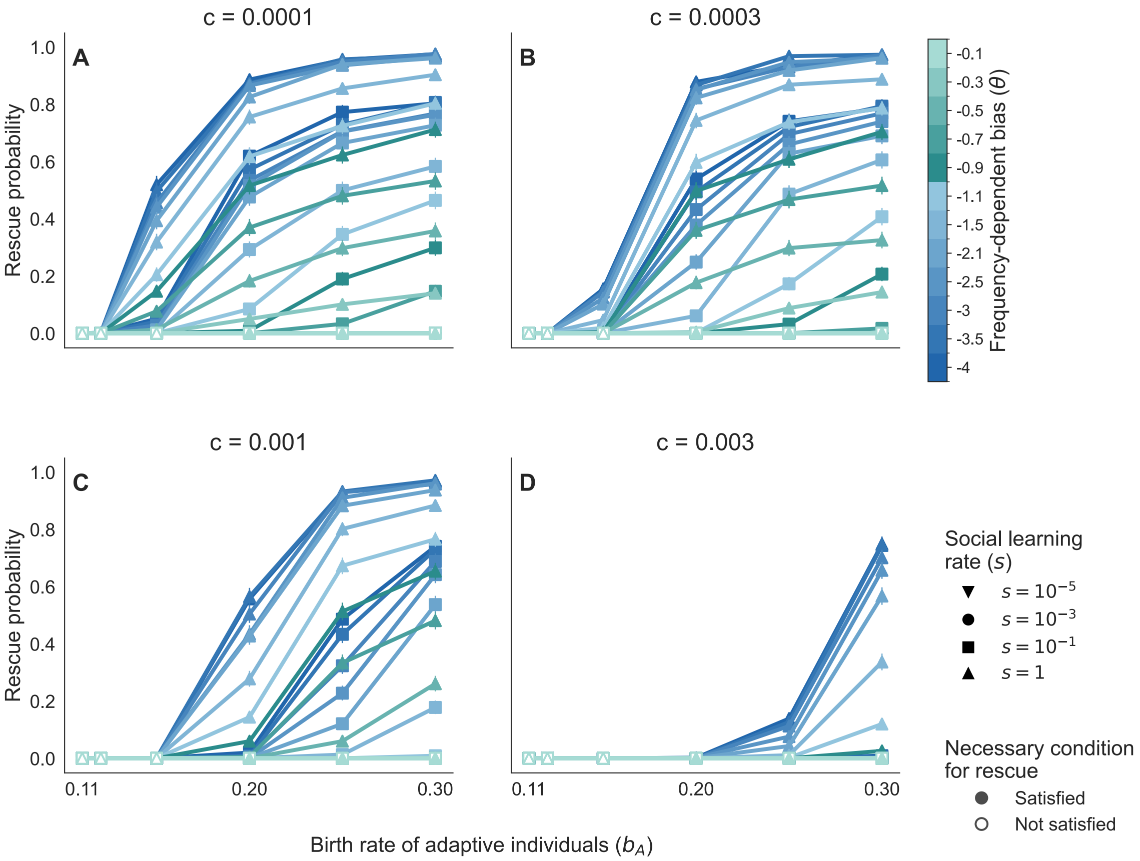

Figure S15: Anticonformity bias promoted cultural evolutionary rescue with a small initial population size Similar to Fig. 5, but the initial population sizes are $N_A(0) = 1$, and $N_M(0) = 9$.

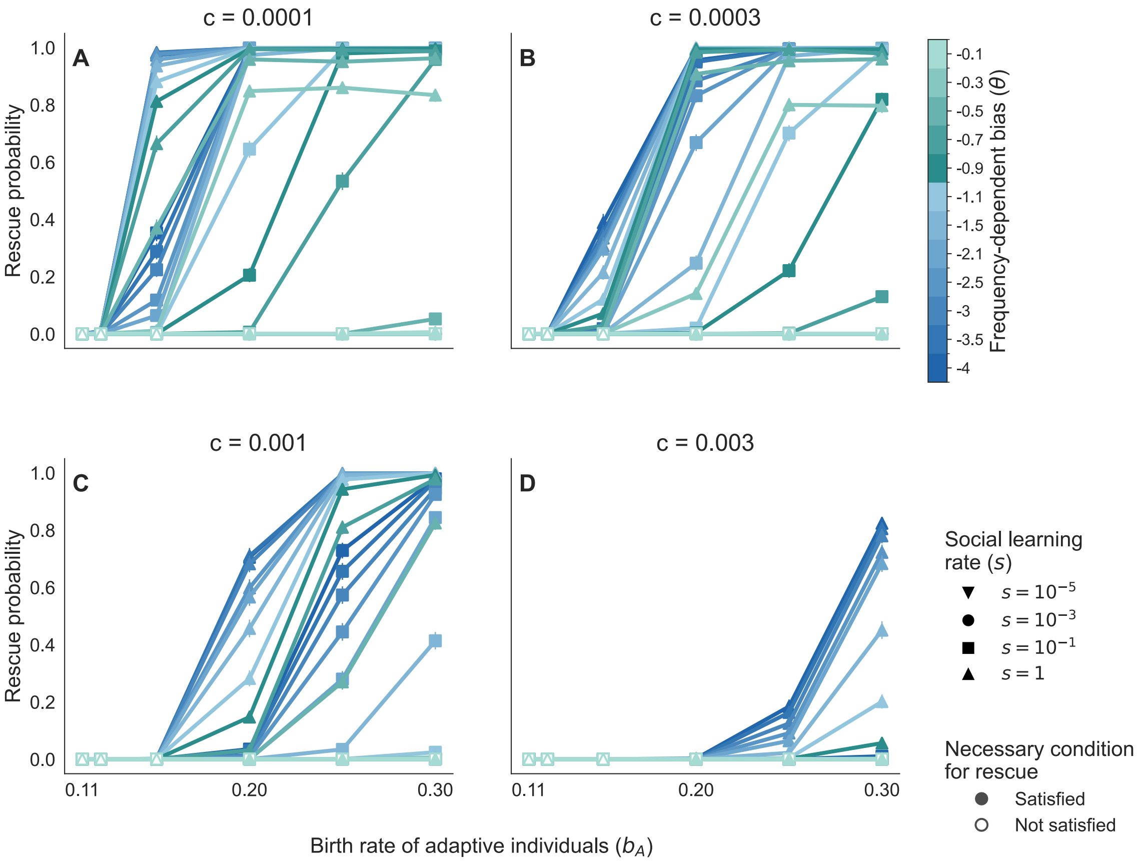

Figure S16: Anticonformity bias promoted cultural evolutionary rescue with a large initial population size

Similar to Fig. 5, but the initial population sizes are $N_A(0) = 1$, and $N_M(0) = 999$.

### S9 Supplementary figures for comparison of rescue probabilities

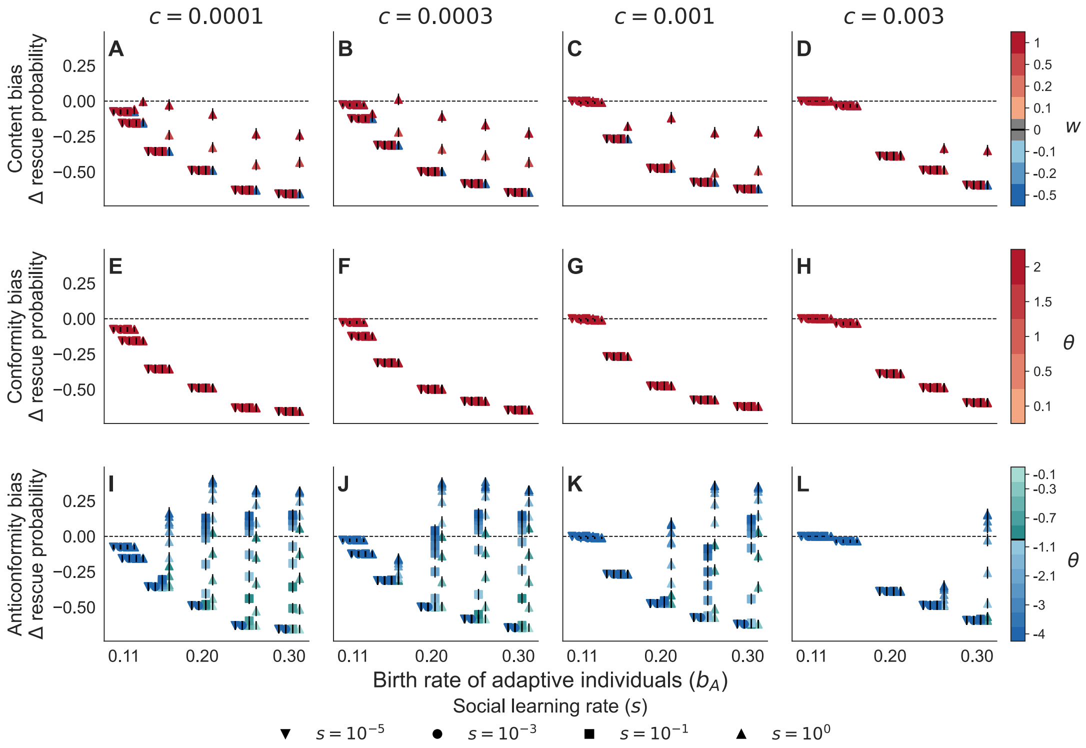

Figure S17: Transmission bias alters when cultural evolutionary rescue outperforms biological evolutionary rescue

Similar to Fig. 6 but $N_M(0) = 9$.

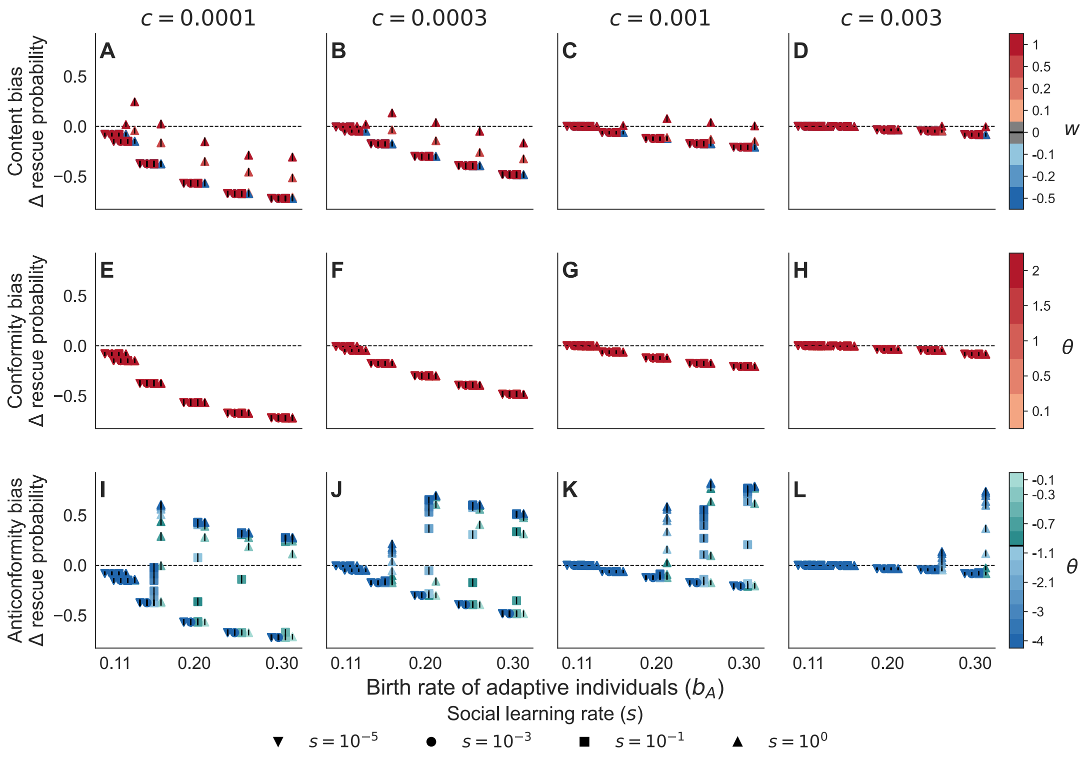

Figure S18: Transmission bias alters when cultural evolutionary rescue outperforms biological evolutionary rescue

Similar to Fig. 6 but $N_M(0) = 999$.

### S10 Supplementary figure and table for rescue probability and establishment time

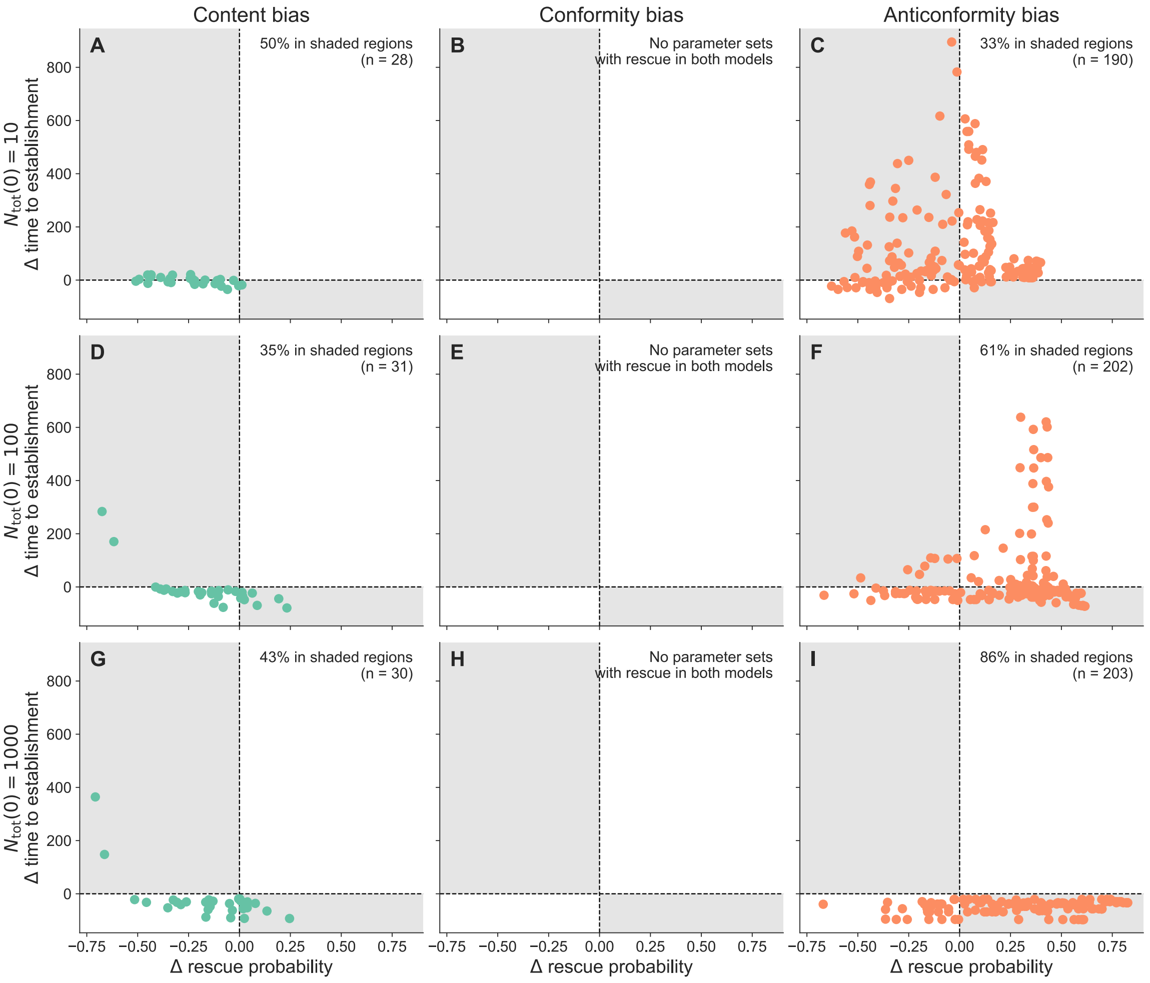

Figure S19: Differences in rescue probability and mean establishment time

Scatter plots showing the relationship between the difference in rescue probability and the difference in conditional mean establishment time (cultural evolution minus biological evolution). Each dot represents a unique combination of parameter values ($b_A$, $c$, $s$, $N_M(0)$, and the strength of transmission bias). The figure removed the cases where rescue probability under either biological or cultural evolution was equal to or lower than 0.01 because establishment time was unreliable there (see Table S1). The shaded areas represent where cultural evolution rescued the population more (less) often and established the adaptive trait faster (slower) than biological evolution. Each row corresponds to a different initial population size (A – C: $N_{\text{tot}} = 10$, D – F: $N_{\text{tot}} = 100$, and G – I: $N_{\text{tot}} = 1000$ ), and each column corresponds to a different transmission bias (A, D, and G: content bias; B, E, H: conformity bias; and C, F, I: anticonformity bias). Parameter values: $\mu = 10^{-3}$ and $l = 0$.

Table S1: Number of parameter sets with rescue probability higher than 0.01

| $N_{\text{tot}}$ | Bias | Both | Biological only | Cultural only | Neither | Total |
|---|---|---|---|---|---|---|
| 10 | Content bias | 28 | 612 | 0 | 128 | 768 |
| 10 | Conformity bias | 0 | 400 | 0 | 80 | 480 |
| 10 | Anticonformity bias | 190 | 770 | 0 | 192 | 1152 |
| 100 | Content bias | 31 | 609 | 0 | 128 | 768 |
| 100 | Conformity bias | 0 | 400 | 0 | 80 | 480 |
| 100 | Anticonformity bias | 202 | 758 | 0 | 192 | 1152 |
| 1000 | Content bias | 30 | 546 | 0 | 192 | 768 |
| 1000 | Conformity bias | 0 | 360 | 0 | 120 | 480 |
| 1000 | Anticonformity bias | 203 | 661 | 0 | 288 | 1152 |
